# Enhancing RNA Delivery: Practical Insights into NeoLNP™ Transfection Reagent

**DOI:** 10.1101/2024.12.10.627676

**Authors:** Xiaobao Chen, Yan Wu, Li Liu, Wei Wang

**Affiliations:** Scindy Pharmaceutical, Co., Ltd., Suzhou Industrial Park, 215125 Suzhou, China; Department of Chemistry, University of Bergen, 5007 Bergen, Norway; Center for Pharmacy, University of Bergen, 5020 Bergen, Norway

**Author notes:** **Corresponding authors:** Xiaobao Chen, Wei Wang.

## Abstract

The rapid advancement of RNA-based therapeutics, particularly in the wake of COVID-19 vaccine success, has prompted significant research into optimizing RNA delivery mechanisms. This study evaluates the NeoLNP™ RNA Transfection Kit developed by Scindy Pharmaceutical, which utilizes lipid nanoparticles (LNPs) for efficient RNA encapsulation and delivery. We systematically investigate various parameters affecting transfection efficiency, including RNA concentration, RNA/LNP volume ratios, mixing techniques, LNP stability, and culture media. Our results demonstrate that the optimal RNA concentration for transfection efficiency is around 40-60 ng/µL, with a 1:0.75-1:1 RNA-to-LNP ratio yielding the highest protein expression. Additionally, we find that gentle mixing techniques outperform harsher methods, and the stability of LNP-RNA complexes significantly influences transfection outcomes. This research provides practical guidelines for enhancing RNA transfection efficiency, paving the way for more effective RNA therapeutics.

## 1. Introduction

The successful deployment of COVID-19 vaccines has significantly accelerated research into RNA technologies, including mRNA, siRNA, saRNA, and others.^1^ RNA holds promising potential as a therapeutic agent in vaccine development,^2^ protein replacement therapies,^3^ and cell therapy applications.^4^ This surge in RNA-related applications has correspondingly led to a marked increase in publications within this field. A key consideration across these applications is ensuring robust expression of RNA in targeted cells. However, two primary challenges must be addressed to achieve high expression levels: RNA sequence design and the choice of delivery vector.^5^ Unlike DNA, RNA molecules are particularly vulnerable to degradation by ribonucleases (RNases). Consequently, prior to transfection, RNA is commonly encapsulated in various transfection reagents, such as cationic polymers,^6^ cationic proteins,^7^ liposomes,^8^ cuboplexes^9^ and lipid nanoparticles (LNPs).^10^ Encapsulated RNA exhibits enhanced stability, transfection efficiency, and expression when compared to naked RNA. Among these vectors, lipid nanoparticles have emerged as the most effective and safe methods for both in vivo and in vitro applications, as evidenced by their successful use in products such as Patisiran,^11^ Comirnaty,^12^ and Spikevax.^13^

Typical formulation of LNP is composed of four main components, ionizable lipid, helper lipid, pegylated lipid and cholesterol, and the ratio of the four lipids is about 50:10:1.5:38.5.^14^ Ionizable lipids are designed to be neutral at physiological pH but become positively charged in acidic environments (such as endosomes).^15^ This charge switch allows them to encapsulate nucleic acids efficiently during formulation by binding to their negatively charged phosphate groups and facilitate endosomal escape after the LNP is internalized by cells. The ionizable lipids destabilize the endosomal membrane, allowing the release of the cargo (e.g., mRNA or siRNA) into the cytoplasm. The helper lipids (e.g., phospholipids like DSPC) help to provide structural integrity and stability to the lipid nanoparticle.^16^ They contribute to membrane fusion and fluidity, aiding the incorporation of various lipids and facilitating delivery and stability of the nanoparticle in biological fluids and during storage. PEGylated lipids have polyethylene glycol (PEG) chains attached to the lipid molecules.^17^ PEG chains reduce opsonization and recognition by the immune system, prolonging the circulation time of the LNPs in the bloodstream, and reduce interactions between LNPs, preventing them from clumping together. Most importantly, they contribute to the uniform size of LNPs by reducing non-specific interactions between the particles. Cholesterol is a stabilizing agent that enhances the fluidity and mechanical strength of the lipid bilayer.^18^ Cholesterol modulates the lipid bilayer’s fluidity and rigidity, ensuring optimal structure. It helps to stabilize the overall structure of the LNP, contributing to its long-term stability and efficient drug delivery. Cholesterol also aids in the fusion of LNPs with the cell membrane, improving cellular uptake.

The encapsulation of RNA with LNPs involves several critical steps in pharmaceutical production.^19^ Due to the solubility constraints of lipids, they are typically dissolved in ethanol (or other organic solvents) and mixed with an aqueous RNA solution using a microfluidic apparatus. This microfluidic mixing process results in a homogeneous size distribution and a low polydispersity index (PDI, often < 0.3).^20^ However, the resulting product contains approximately 25% ethanol, making it unsuitable for immediate use in RNA transfection experiments or *in vivo* studies. To address this, ethanol (or other organic solvents) is removed via dilution, ultrafiltration, or dialysis in laboratory settings, while tangential flow filtration (TFF) is commonly used in pharmaceutical manufacturing.^21^ Buffer exchange can also occur during this step, ensuring that the LNPs transition to a biocompatible, aqueous environment, suitable for therapeutic applications. To safeguard against bacterial contamination, LNPs are typically sterilized through filtration (e.g., 0.22 µm filters). The entire process can take days to complete, significantly slowing down research and development in RNA transfection and related studies.^22^

In response to these challenges, Scindy Pharmaceutical has developed the NeoLNP™ RNA Transfection Kit, designed to streamline RNA encapsulation via LNP technology, accelerating the process and removing common bottlenecks in RNA research. The NeoLNP™ RNA Transfection Kit is a ready-to-use LNP solution. Its operation couldn’t be simpler—just mix the kit with the RNA solution, and encapsulation occurs spontaneously and simultaneously during the mixing. Once mixed, the RNA is securely encapsulated within the LNPs, ready for immediate use in the next phase of research. In this study, we investigate the influence of various factors (such as: RNA concentrations, N/P ratio, mixing method, stability of RNA LNP, culture media, RNA types, etc.) on transfection efficiency, offering practical guidelines for the optimal application of the NeoLNP™ RNA Transfection Kit.

## 2. Materials and Protocols

NeoLNP Transfection Kit (REF: SDR8006) was obtained from Scindy Pharmaceuticals. The kit contains a transfection reagent and buffer solution for prepare RNA-LNP complex. The cell lines used in this study included HEK 293T, A549, Jurkat, and THP-1. Cells were cultured at 37 °C with 5% CO_2_. Details of the culture medium, seeding density and commercial sources of cells are shown in Table 1. The EGFP mRNA (cat. no. DD4503-02) and Luc mRNA (cat. no. DD4511-02) were purchased from Vazyme Biotech Co., Ltd. The saRNA (cat. no. HBPM00011-1) was purchased from Hzymes Biotchnology Co., Ltd. and circRNA (cat. no. HZ010) was obtained from CATUG biotechnology. Reporter gene expression levels, cell viability, and protein concentration were determined using fluorescent microscopy (Keyence BZ-X800) and flow cytometry (Beckman Coulter, Cytoflex S).

**Table 1.**
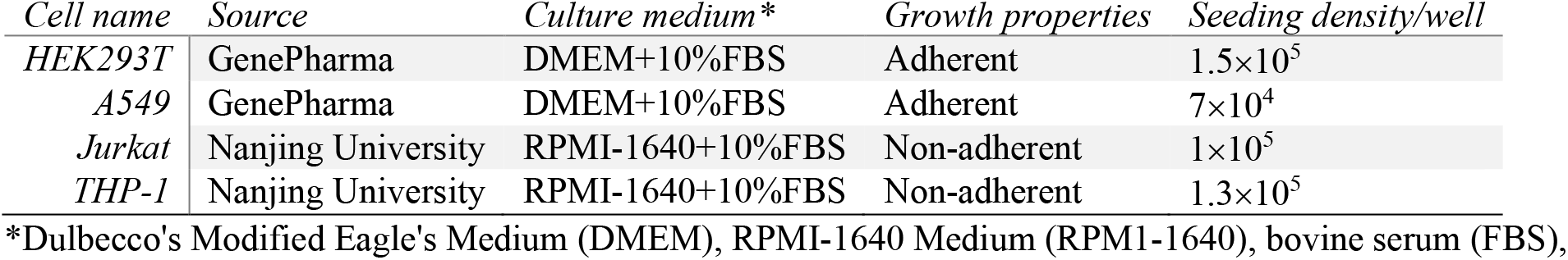
Cell lines, commercial sources, culture medium, and seeding density.

| *Cell name* | *Source* | *Culture medium\** | *Growth properties* | *Seeding density/well* |
| --- | --- | --- | --- | --- |
| *HEK293T* | GenePharma | DMEM+10%FBS | Adherent | $1.5 \times 10^5$ |
| *A549* | GenePharma | DMEM+10%FBS | Adherent | $7 \times 10^4$ |
| *Jurkat* | Nanjing University | RPMI-1640+10%FBS | Non-adherent | $1 \times 10^5$ |
| *THP-1* | Nanjing University | RPMI-1640+10%FBS | Non-adherent | $1.3 \times 10^5$ |

\*Dulbecco's Modified Eagle's Medium (DMEM), RPMI-1640 Medium (RPMI-1640), bovine serum (FBS),

Unless otherwise noted, RNA transfections were performed following the procedure described in the NeoLNP Transfection Kit package insert. Briefly, (1) Cells were cultured overnight in 24-well trays at 100,000 to 150,000 cells per well; (2) the next day, RNA-NeoLNP complexes were freshly prepared (Figure 1) and added directly to the wells; (3) the cells were incubated with the complexes for the recommended time; and (4) expression of reporter gene proteins was assayed at 6 hours post-transfection or when otherwise indicated. RNA-NeoLNP complexes were prepared with a buffer solution provided in the kit to maintain RNA stability. All experiments were done in triplicate to ensure reproducibility. Statisical results are found in the Supplementary Information.

**Figure 1.**
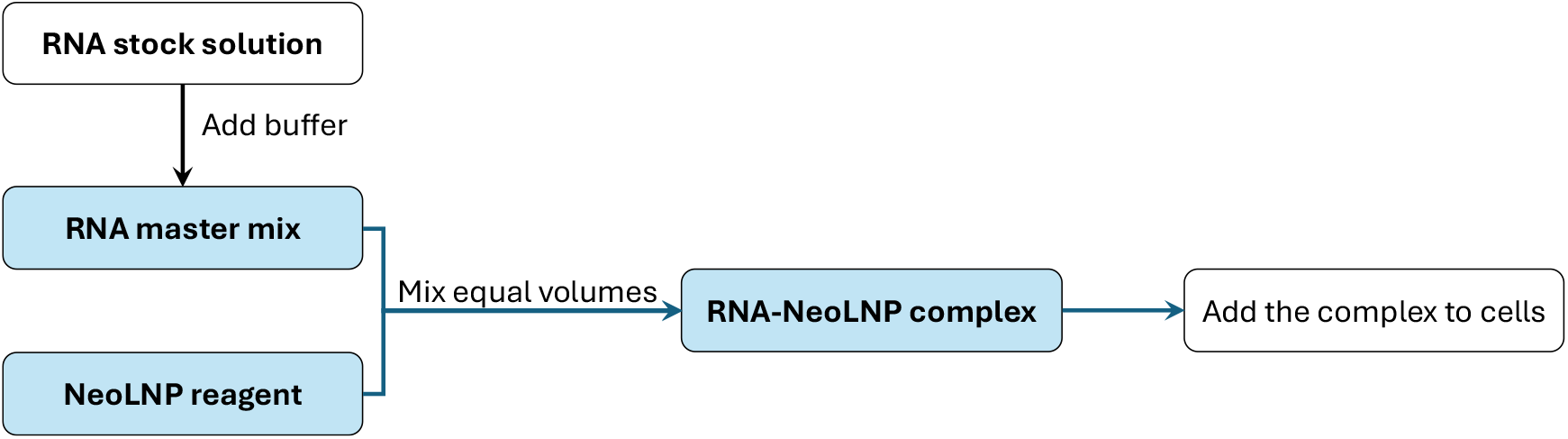
A schematic flow chart for preparation of RNA-NeoLNP complex before transfection experiment.

## 3. Factors for optimal transfection using NeoLNP

### 3.1 The effect of mRNA concentration on the transfection

We investigated the impact of mRNA concentration on transfection efficiency by preparing LNP-mRNA complexes. A 1 µg/µL EGFP-mRNA stock solution was diluted to final concentrations of 20, 40, 60, 80, and 100 ng/µL using the provided buffer, while a 0.1 µg/µL NeoLNP solution was similarly diluted to match these concentrations. Equal volumes of the mRNA and LNP solutions were combined at a 1:1 ratio to form LNP-mRNA complexes, which were introduced into 293T cells at a dose of 100 ng per well. Transfection efficiency was assessed using microscopy and flow cytometry. Microscopy enabled visual confirmation of reporter protein expression, while flow cytometry provided quantitative measurements of protein expression across the range of mRNA concentrations. By comparing these data, we aimed to establish any correlations between mRNA concentration and transfection efficiency. This analysis is critical for optimizing mRNA delivery systems by demonstrating the key role of mRNA concentration in effective transfection.

The results (Figure 2, Table S1 and S2) showed that, compared to the blank control, which exhibited a relative mean flourescence intensity (RFL) of 556, the introduction of LNP-mRNA complexes significantly enhanced protein expression at all concentrations. Notably, the highest protein expression was observed at 60 ng/µL, with an average value of 1.72E+06, followed closely by the 80 ng/µL concentration at 1.71E+06. Although the expression slightly decreased at the 100 ng/µL concentration (1.57E+06), the overall trend indicated an increase in transfection efficiency with mRNA concentration up to 60 ng/µL. The standard deviations were relatively low, suggesting consistent results across the replicates, particularly at lower concentrations. These findings highlight the importance of optimizing mRNA concentration for effective transfection using LNPs. However, the differnece of expression efficience is not great influnced by the mRNA concentration from 20 ng/µL to 100 ng/µL.

**Figure 2.**
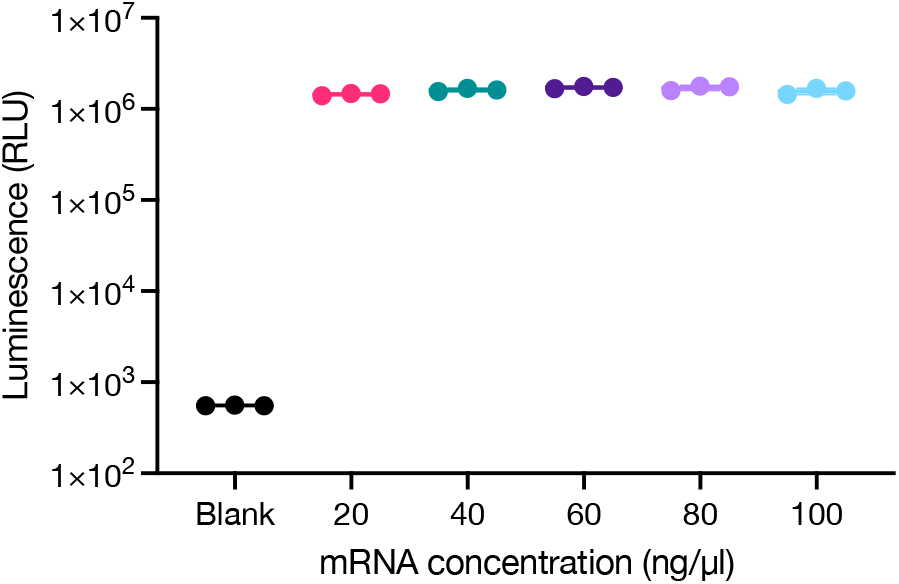
The fluoresense intensity changes with different mRNA concentration the volume ratio of mRNA/NeoLNP is 1:1.

### 3.2 The effect of mRNA per well on the transfection

In our experiment assessing the effect of LNP concentration on transfection efficiency, we prepared LNP-mRNA complexes by diluting a 1 µg/µL Luc-mRNA solution to 40 ng/µL and similarly diluting a 0.1 µg/µL LNP solution to the same concentration. The two solutions were combined at two volume ratios (1:1 and 1:0.5) and thoroughly mixed by pipetting at least 20 times to form the LNP-mRNA complex. We then introduced varying doses of the complex—50 ng, 100 ng, and 200 ng per well—into the cells to evaluate the relationship between LNP concentration and protein expression. We anticipate that higher doses of the LNP-mRNA complex will yield increased protein expression, but the results showed difference.

For the 293T cell line, at a mRNA/LNP ratio of 1:0.5, the average protein expression increased with mRNA dose, peaking at 200 ng/well with a RFL of 1.23E+06. In contrast, at a mRNA/LNP ratio of 1:1, the average expression decreased with increasing mRNA concentration, with the highest value observed at 50 ng/well (RFL=1.09E+06) and a notable drop to 6.64E+05 at 200 ng/well. The blank control averaged 5.51E+02, indicating a significant enhancement in transfection efficiency with LNP-mRNA complexes.

For the A549 cell line, similar trends were observed. At a 1:0.5 ratio, average protein expression also peaked at 200 ng/well, reaching RFL:1.34E+06. Conversely, the 1:1 ratio showed a decrease in expression at higher doses, with an average of 1.14E+06 at 50 ng/well and dropping to 1.04E+06 at 200 ng/well. The blank control averaged 5.15E+02. These results indicate that both the mRNA concentration and the ratio of mRNA to LNP significantly influence transfection efficiency, emphasizing the importance of optimizing these parameters for effective mRNA delivery in different cell lines.

**Figure 3.**
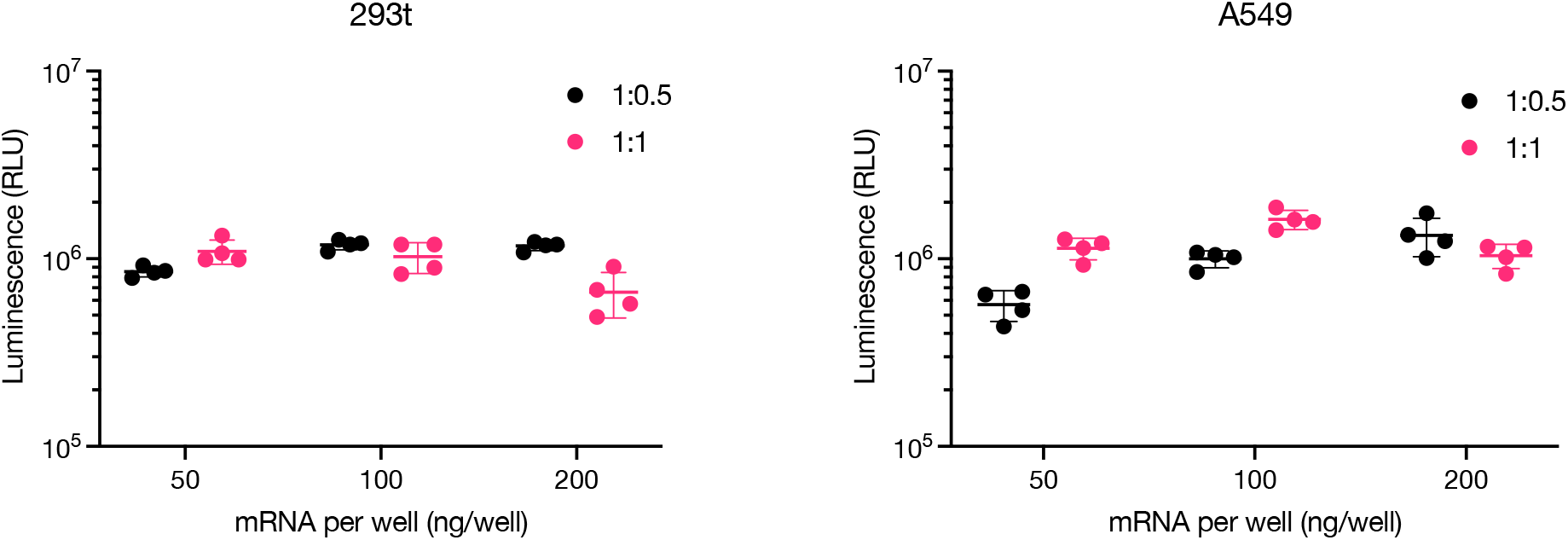
The fluoresense intensity changes with adding different mRNA per well on two cell lines at two mRNA/NeoLNP ratio.

### 3.3 The effect of mRNA and LNP charge ratio on transfection

The effect of varying mRNA and LNP ratios on transfection efficiency was studied. To prepare the working solutions, we diluted a 1 µg/µL EGFP-mRNA stock to a concentration of 40 ng/µL using buffer. Simultaneously, the 0.1 µg/µL LNP solution was diluted stepwise to final concentrations of 20, 30, 40, 50, and 60 ng/µL, also using the same buffer. Equal volumes of the diluted mRNA solutions were combined with the 40 ng/µL LNP solution at a 1:1 ratio through pipetting to create the respective LNP-mRNA complexes. These complexes were introduced into 293T cells at a final concentration of 100 ng per well to evaluate protein expression levels. The assessment of transfection efficiency was conducted using both fluorecent microscopy and flow cytometry. We expect that the varying LNP concentrations will impact the delivery and expression of mRNA in the cells, providing insights into optimizing the mRNA-LNP formulation for enhanced transfection outcomes.

The results demonstrated a clear correlation between the mRNA to LNP ratio and protein expression levels. The optimal ratio appeared to be 1:0.75, achieving the highest average expression of RFL:1.74E+06, significantly surpassing the blank control average of 556. As the ratio increased to 1:1.25 and 1:1.5, there was a noticeable decline in mean fluorescence intensity, dropping to 1.43E+06 and 1.08E+06, respectively. The standard deviations were relatively low, indicating consistent results across replicates, particularly at the lower ratios. These findings suggest that while the electrostatic interactions between the negatively charged mRNA and positively charged LNP are crucial for effective complex formation, an excess of LNP beyond the optimal ratio may hinder transfection efficiency.

**Figure 4.**
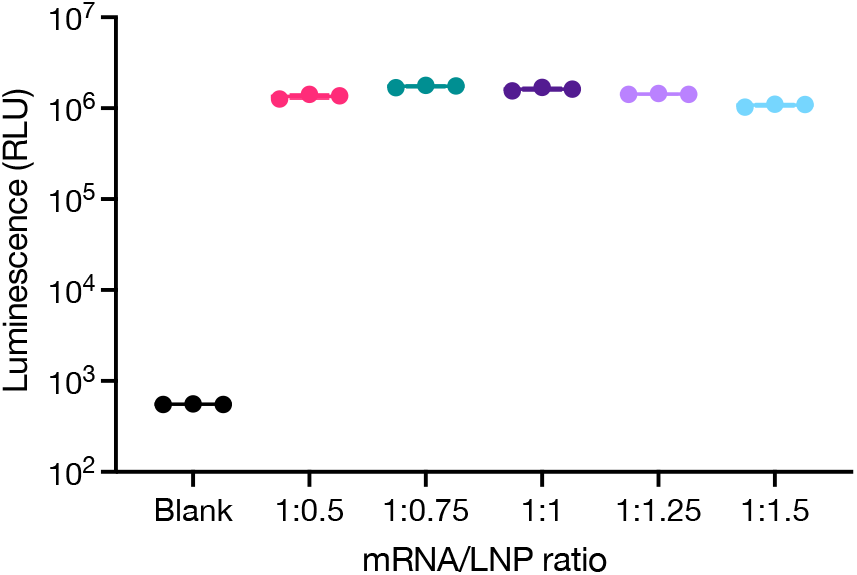
The mRNA-NeoLNP transfection efficiency affected by the mixing ratio of mRNA and NeoLNP. Five different volume ratios were studied.

### 3.4 Mixing techniques

In our study on the impact of mixing techniques on the transfection efficiency of mRNA-LNP complexes, we utilized flow cytometry to analyze two mRNA/NeoLNP ratios: 1:1 and 1:0.5, using three different mixing methods: repetitive pipetting, vortexing for 3 seconds, and vortexing for 10 seconds. For the 1:1 ratio, the average protein expression was highest with repetitive pipetting (1.70E+06), followed closely by vortexing for 3 seconds (1.64E+06) and then vortexing for 10 seconds (1.56E+06). However, the difference is insignificant. The relatively low standard deviations, particularly for the pipetting method, indicate consistent results, suggesting that pipetting may facilitate more effective mixing of mRNA and LNP, optimizing complex formation.

In contrast, at the 1:0.5 ratio, the transfection efficiencies were significantly lower across all mixing methods, with average protein expressions ranging from 4.41E+05 to 4.62E+05, compared to the 1:1 ratio. The blank control averaged 2197, underscoring the effect of LNP concentration on transfection outcomes. Standard deviations were also low for this ratio, particularly for the vortexing for 10 seconds method, indicating that the mixing method had less influence on the transfection efficiency at this lower ratio.

These results highlight the importance of optimizing both the mRNA to LNP ratio and the mixing technique to maximize transfection efficiency. The superior performance of repetitive pipetting at the 1:1 ratio suggests that gentle mixing may promote better interaction between mRNA and LNP, whereas the lower efficiencies at the 1:0.5 ratio indicate that insufficient LNP concentration may limit transfection success, regardless of mixing method.

**Figure 5.**
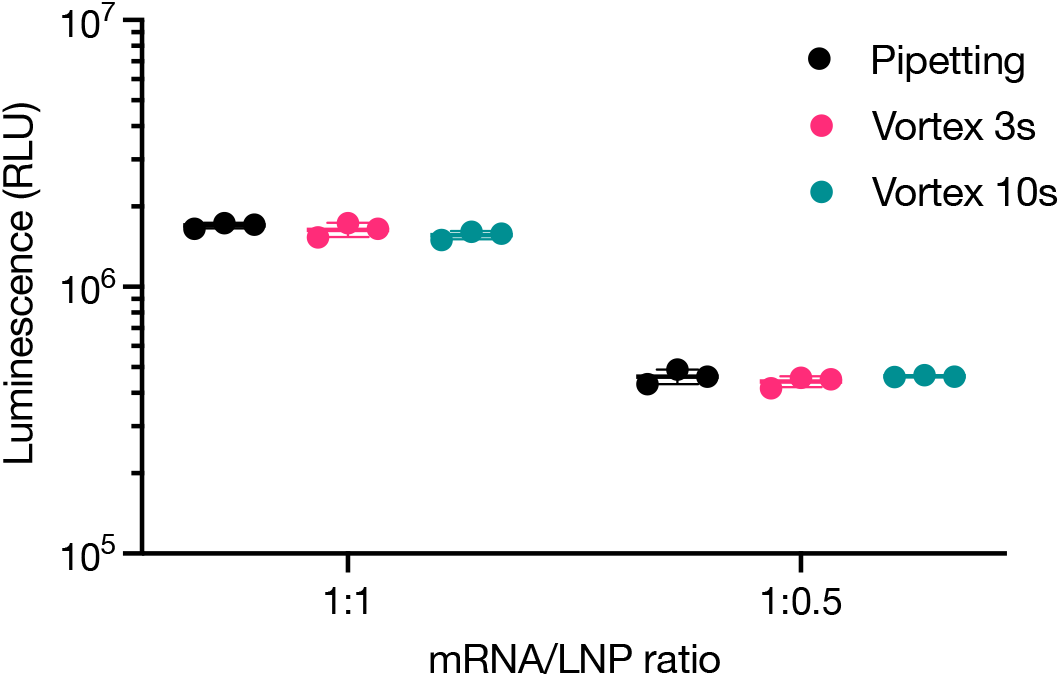
The affect of mxing techniques on the mRNA-NeoLNP transfection. Three mixing techniques were tested, with pepetting up-and-down for 20 times, vortexing for 3 seconds and vortexing for 10 seconds.

### 3.5 mRNA-LNP complex stability

In our investigation of the stability of mRNA-LNP complexes, we formulated EGFP-mRNA and LNP at a concentration of 40 ng/µL. The LNP-mRNA complexes were prepared through simple pipetting. After forming the complexes, they were incubated at room temperature for varying intervals of 0, 0.5, 1, 3, 8, and 24 hours.

During this time points, the mRNA-LNP complexes were placed in the room temperature. Following incubation, the complexes were introduced into 293T cells at a dose of 100 ng per well to assess fluorescence intensity.

For the 1:1 ratio, the average protein expression increased from 1.62E+06 at time zero, reaching a peak of 2.89E+06 at 24 hours. The initial rise in expression was consistent across the first 0.5 hours, with minor fluctuations, suggesting that the complexes remained stable and functional during this period. The standard deviations were low, particularly at the earlier time points, indicating reliable results.

For the 1:0.5 ratio, the average expression also increased from 1.71E+06 at time zero to 3.02E+06 at 24 hours, peaking at 3.02E+06. The data demonstrated a consistent upward trend, particularly notable at the 1-hour mark, where the average expression reached 3.02E+06, suggesting that this ratio might provide improved stability and transfection efficiency compared to the 1:1 ratio. However, the standard deviations at certain time points were higher, indicating more variability in the results, particularly at the later time points.

These results highlight the importance of the stability of mRNA-LNP complexes in transfection efficiency. The increasing trends in protein expression over time for both ratios indicate that these complexes maintain their functionality over extended periods. However, the more consistent performance at the 1:1 ratio suggests that it might be more suitable for applications requiring immediate results, while the 1:0.5 ratio shows promise for sustained expression.

**Figure 6.**
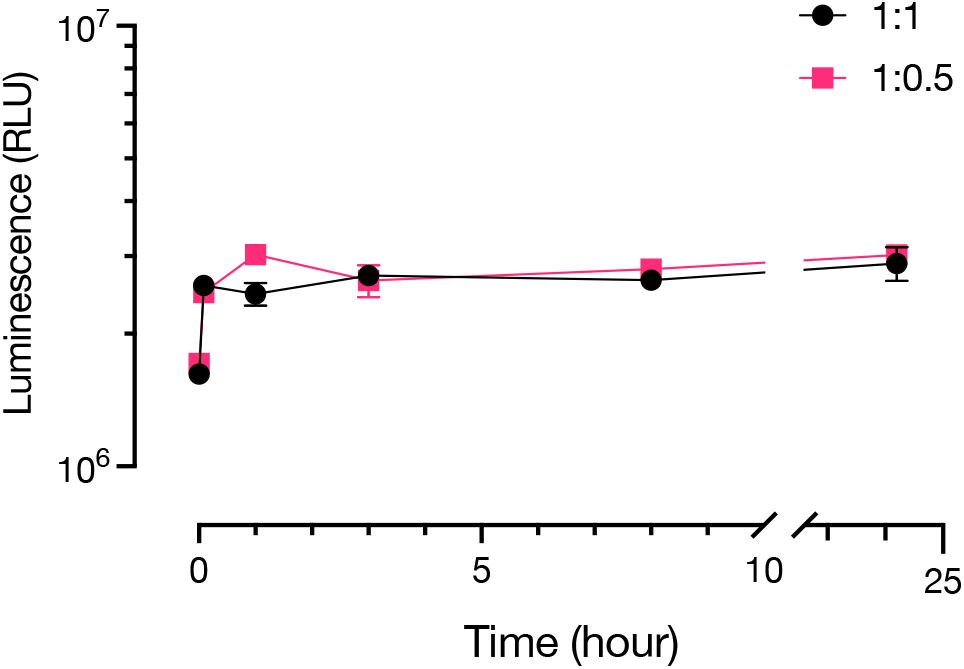
The stability of mRNA-NeoLNP complex stored at room temperature upto 24 hours. The transfection efficiency is not significantly affected.

### 3.6 The effect of culture medium on the RNA transfection

In our study of the influence of culture medium on transfection efficiency, we prepared EGFP-mRNA and LNP at 40 ng/µL, forming the LNP-mRNA complexes through pipetting to mix mRNA with LNP. These complexes were then introduced into three different culture media conditions: serum(+)/antibiotics(-), serum(-)/antibiotics(-), and serum(+)/antibiotics(+). The results were analyzed for two mRNA to LNP ratios: 1:1 and 1:0.5.

For the 1:1 ratio, the average protein expression was highest in the serum(-)/antibiotics(-) condition (1.93E+06), followed by serum(+)/antibiotics(-) (1.70E+06), and lowest in the serum(+)/antibiotics(+) condition (1.64E+06). The standard deviations were relatively low, particularly for the serum(-)/antibiotics(-) condition, indicating reliable data across replicates. These findings suggest that the culture media may have limited effect on the transfection efficiency in this ratio, while the addition of serum and antibiotics might hinder the expression levels.

In contrast, for the 1:0.5 ratio, the average protein expressions were lower overall, with the highest average seen in the serum(-)/antibiotics(-) condition (6.63E+05) and the lowest in the serum(+)/antibiotics(+) condition (4.56E+05). The standard deviations for the three condition were also low, indicating consistency in the data. The lower expression levels across all conditions compared to the 1:1 ratio suggest that the mRNA concentration may be a limiting factor for transfection efficiency at this ratio. Overall, these results emphasize the robust transfection efficiency at all culture media, with or without serum and antibiotics.

**Figure 7.**
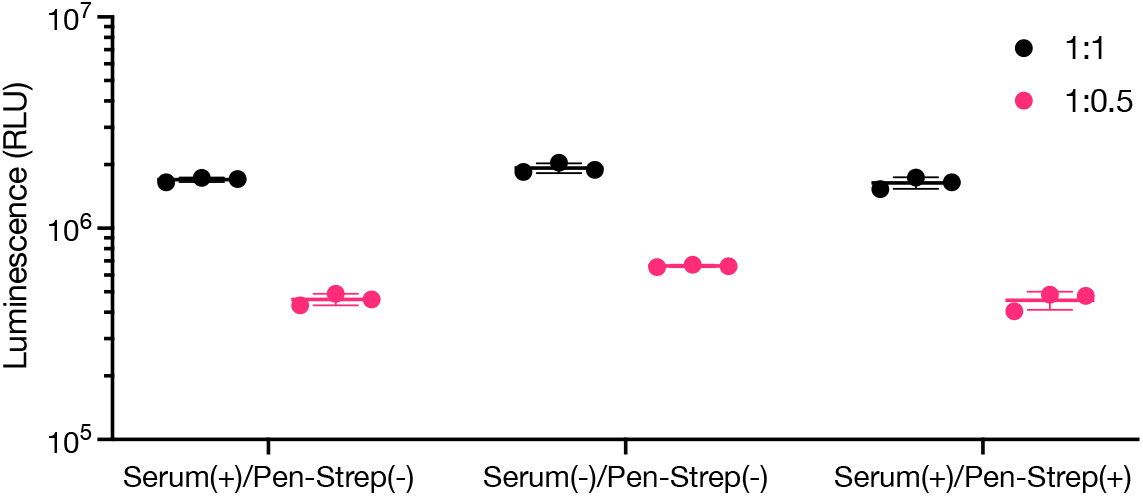
The flourescence intensity of EGFP mRNA transfection on 293t cells with two mRNA-to-NeoLNP ratios (1:1 and 1:0.5) in three differen culture media with or without serum and antibotics.

### 3.7 The change of transfection intensity with time

In the experiment investigating the dynamics of fluorescence intensity over time after transfection with Luc-mRNA, we prepared both Luc-mRNA and LNP at a concentration of 40 ng/µL. The LNP-mRNA complexes were formed through pipetting and mixing, and these complexes were introduced into two cell lines, 293T and A549, at a concentration of 100 ng per well. Fluorescence intensity was monitored at multiple time points: 6 h, 12 h, 24 h, 48 h, 72 h, and day 7.

For the 293T cells, the average fluorescence intensity for the mRNA to LNP ratio of 1:0.5 peaked at 12 hours (1.26E+06), then declined over the following time points, with a notable drop by 72 hours (1.71E+05). The blank control consistently showed low fluorescence, confirming the specific expression from the transfected cells. The 1:1 ratio also exhibited a peak fluorescence intensity at 6 hours (6.97E+05), but this value diminished significantly by the 48-hour mark (4.43E+05).

For the A549 cells, the 1:0.5 ratio showed a peak average fluorescence intensity of 1.00E+06 at 24 hours before decreasing to 4.24E+05 by 72 hours. In contrast, the 1:1 ratio had its maximum fluorescence (1.62E+06) at the 24-hour time point, followed by a decline to 3.98E+05 at 48 hours. The LipoMax formulation outperformed both ratios at all time points, reaching a peak of 1.98E+06 at 24 hours, but subsequently dropped to 2.22E+05 by 72 hours.

These results suggest that the transfection efficiency is time-dependent, with an optimal protein expression observed at earlier time points. The decline in fluorescence intensity over time may indicate that the stability of the mRNA-LNP complexes decreases, leading to a reduction in transgene expression. However, mRNA sequece is also an important factor on the stability of mRNA in cytoplasma. RNA sequence design is a important stabilization strategies for mRNA-LNP complexes to prolong protein expression.

**Figure 8.**
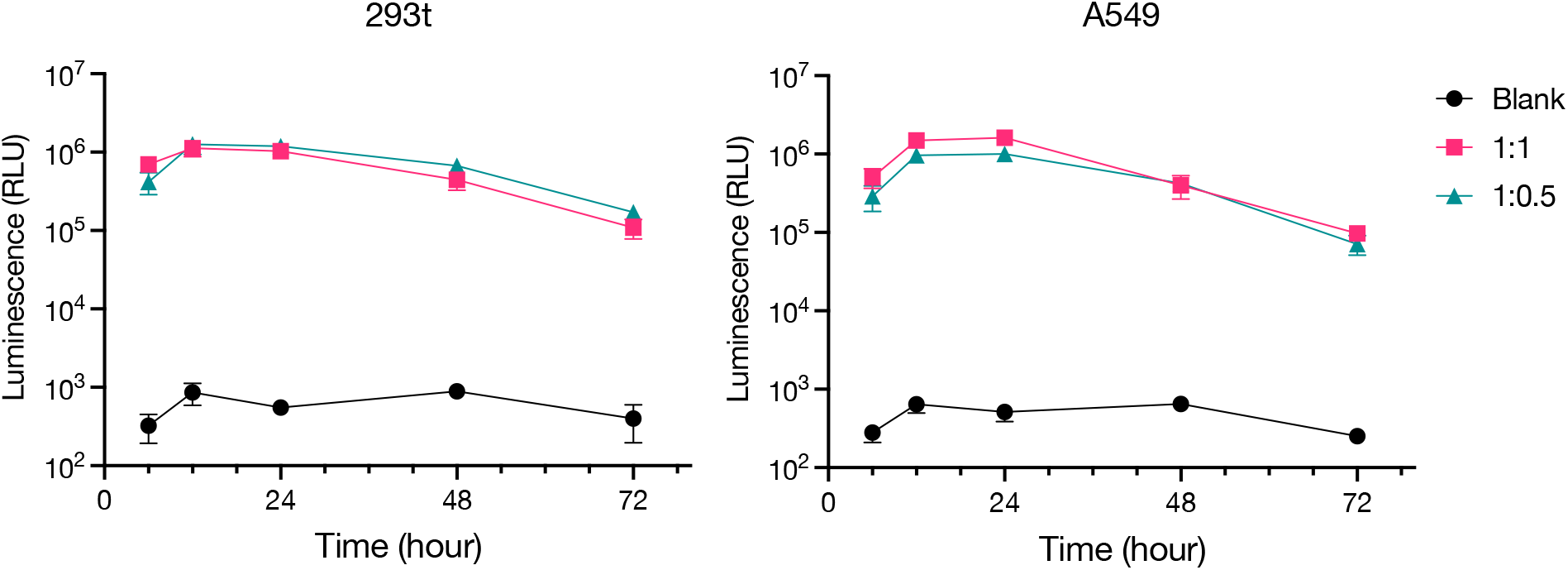
The dynamic flourescence intensity over 72 hours after EGFP-mRNA transfection on 293t and A549 cells with two mRNA-to-NeoLNP ratios (1:1 and 1:0.5).

### 3.8 RNA types

We investigated the transfection efficiency of various RNA types using NeoLNP as the delivery vehicle. The RNA solutions, including circRNA, saRNA, and mRNA, were initially prepared at 1 µg/µL and subsequently diluted to 40 ng/µL using the buffer provided in the kit. The LNP solutions were also diluted to the same concentration. LNP-RNA complexes were formed by mixing these solutions in a 1:1 volume ratio, and the resulting complexes were introduced into 293T cells at a dose of 100 ng per well to assess protein expression through quantitative flow cytometry.

The results demonstrated that at a 1:1 RNA-to-NeoLNP ratio, mRNA achieved the highest average fluorescence intensity of 1.70E+06, indicating effective transfection and robust protein expression. This result aligns with the well-established efficiency of mRNA in transfection applications, showcasing its potential for delivering genetic material into target cells. In comparison, saRNA exhibited a considerable expression level of 1.20E+06, while circRNA showed the lowest expression at 4.96E+05. The reduced efficacy of circRNA may be attributed to its structural characteristics and stability, which can impact its uptake and subsequent translation in the target cells.

When the RNA-to-NeoLNP ratio was adjusted to 1:0.5, a notable decline in average fluorescence intensity was observed for mRNA, dropping to 4.60E+05. This reduction highlights the importance of optimizing the RNA-to-LNP ratio to ensure efficient transfection. In this ratio, saRNA maintained a relatively high expression level of 1.08E+06, indicating its resilience in transfection efficiency despite the lower LNP concentration. Conversely, circRNA remained low at 1.94E+05, reaffirming its limited effectiveness in this context.

These findings suggest that mRNA is the most effective RNA type for transfection when complexed with NeoLNP at a 1:1 ratio, followed by saRNA. The lower transfection efficiency observed at the 1:0.5 ratio underscores the necessity of careful optimization of LNP formulations and RNA types for successful gene delivery.

**Figure 9.**
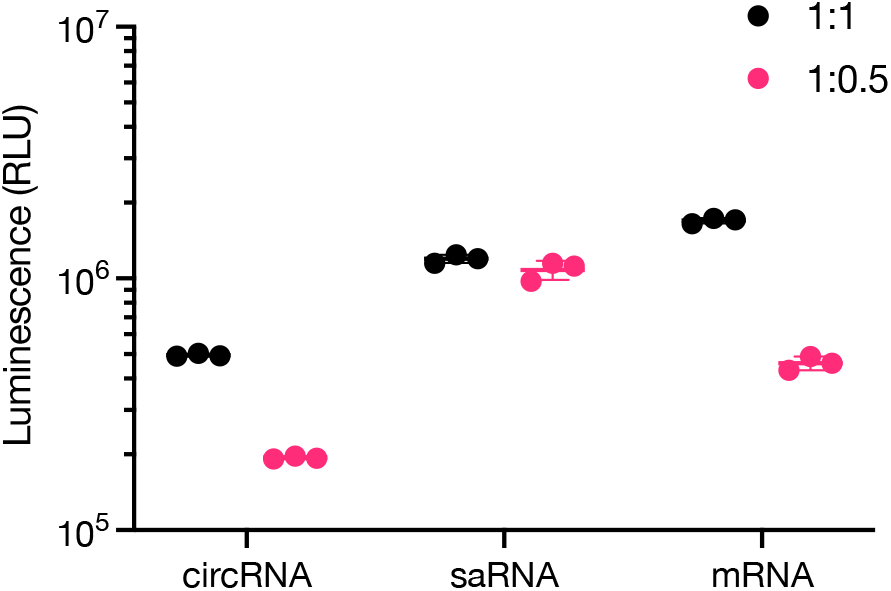
The flourescence intensity of EGFP-circRNA, saRNA, and mRNA transfection on 293t cells with two mRNA-to-NeoLNP ratios (1:1 and 1:0.5).

### 3.9 Transfection efficiency in different cell lines

We investigated the transfection efficiency of EGFP-mRNA delivered via LNPs across different cell lines: 293T, Jurkat, and THP-1. Perticularlly, Jurkat and THP-1 cell lines are known to be be challenging to transfect. Both the mRNA and LNP were prepared at a concentration of 40 ng/µL, and complexes were formed through pipetting. These LNP-mRNA complexes were applied to the cell lines at a dose of 100 ng per well.

At a 1:1 mRNA-to-NeoLNP ratio, the results revealed significant variations in transfection efficiency among the cell lines. The 293T cell line exhibited the highest average fluorescence intensity of 1.70E+06, indicating a robust transfection efficiency and effective protein expression. In contrast, Jurkat cells displayed lower expression levels, with an average fluorescence intensity of only 1.15E+04. The %EGFP+ is about 50% for Jurkat cells. In contrast, %EGFP+ for THP-1 cells is 84% and it for 293t cells is over 99.5%. This stark difference suggests that 293T cells are more amenable to transfection with LNPs than Jurkat cells, potentially due to their higher susceptibility to lipid-mediated transfection methods. THP-1 cells demonstrated an intermediate expression level of 3.03E+05, suggesting a moderate transfection efficiency relative to the other two cell lines.

When the mRNA-to-NeoLNP ratio was adjusted to 1:0.5, the transfection efficiency across all cell lines decreased. 293T cells maintained a respectable average expression level of 4.60E+05, while Jurkat cells saw a further reduction, averaging only 6.34E+03. THP-1 cells also exhibited diminished expression, averaging 9.46E+04. The %EGFP+ is about 10% to 13% for Jurkat cells. In contrast, %EGFP+ for THP-1 cells is 67% and it for 293t cells is over 97%. The substantial drop in fluorescence intensity in Jurkat cells highlights their reduced responsiveness to lower LNP concentrations.

Overall, these findings underscore the importance of the optimization of RNA-to-LNP ratios for effective transfection for differnet cell lines. Additionally, the results emphasize the need to further explore the cellular factors influencing transfection efficiency in Jurkat and THP-1 cells, potentially guiding the development of tailored strategies for enhancing RNA delivery in challenging cell types.

**Figure 10.**
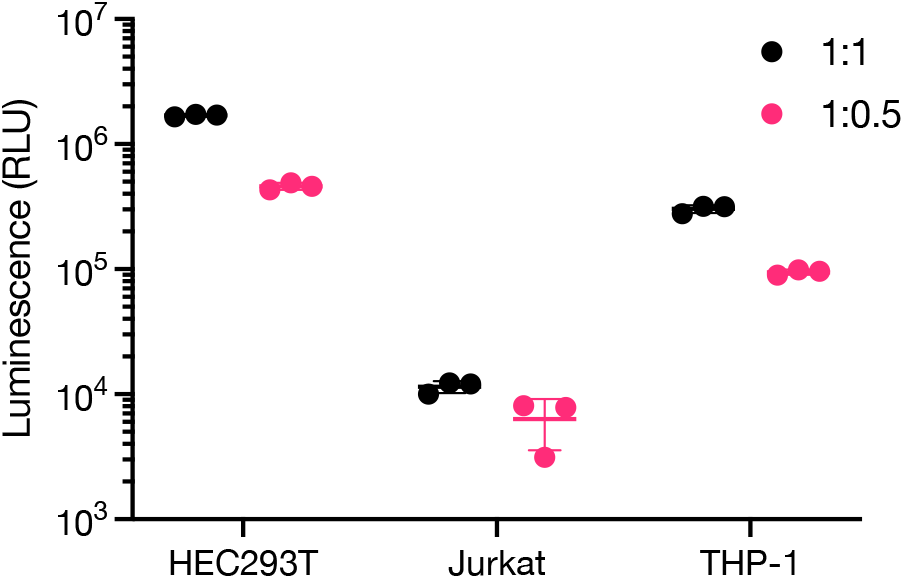
The flourescence intensity of EGFP posttransfection on 293t, Jurkat and THP-1 cell lines with two mRNA-to-NeoLNP ratios (1:1 and 1:0.5).

## 4. Conclusion

In summary, our investigation into the NeoLNP™ RNA Transfection Kit highlights critical factors influencing the efficacy of RNA transfection using lipid nanoparticles. Both mRNA concentration and the mRNA-to-LNP ratio significantly impact transfection efficiency across various cell lines. Additionally, while mixing techniques and culture media play roles, the stability of mRNA-LNP complexes appears to be a crucial determinant of performance. The results underscore that optimization of parameters like RNA type and transfection conditions can drastically enhance gene delivery effectiveness. Furthermore, the findings suggest avenues for future research, particularly in refining strategies for challenging cell types such as Jurkat and THP-1, thereby paving the way for improved therapeutic applications of RNA-based technologies. The NeoLNP™ RNA Transfection Kit stands out as a promising tool for streamlining RNA delivery in research and clinical settings.

