## Supplemental Table 1 for "Enhancing RNA Delivery: Practical Insights into NeoLNP™ Transfection Reagent"

### 1. Effect of EGFP-mRNA and LNP concentration on transfection

**Table S1:** Microscopic images showing the effect of EGFP-mRNA concentration on transfection efficiency at varying doses.

| blank | 20 ng/µl | 40 ng/µl | 60 ng/µl | 80 ng/µl | 100 ng/µl |
|---|---|---|---|---|---|
| 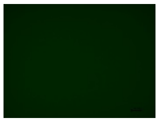 | 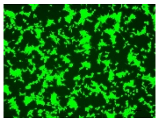 | 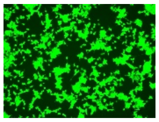 | 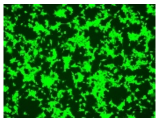 | 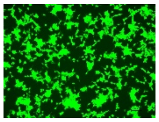 | 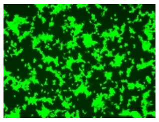 |

**Table S2:** Flow cytometry data showing transfection efficiency across different EGFP-mRNA concentrations.

| | blank | 20 ng/µl | 40 ng/µl | 60 ng/µl | 80 ng/µl | 100 ng/µl |
|---|---|---|---|---|---|---|
| 1 | 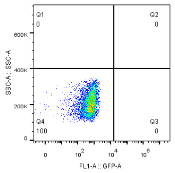 | 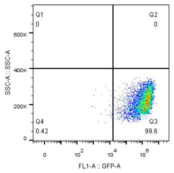 | 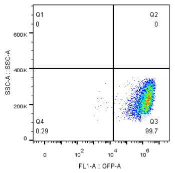 | 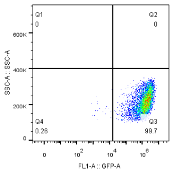 | 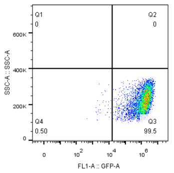 | 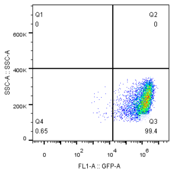 |
| 2 | 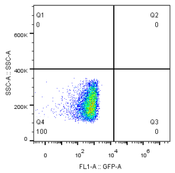 | 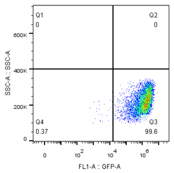 | 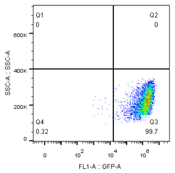 | 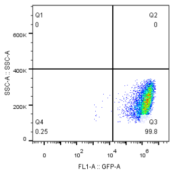 | 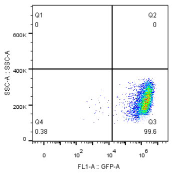 | 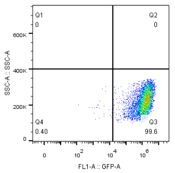 |
| 3 | 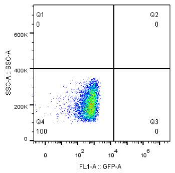 | 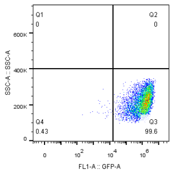 | 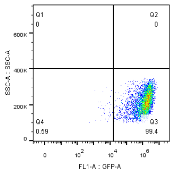 | 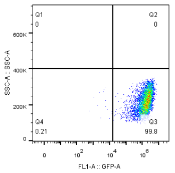 | 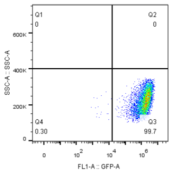 | 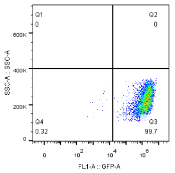 |
| Data 1 | 562 | 1.48E+06 | 1.69E+06 | 1.67E+06 | 1.59E+06 | 1.44E+06 |
| Data 2 | 552 | 1.47E+06 | 1.62E+06 | 1.73E+06 | 1.76E+06 | 1.58E+06 |
| Data 3 | 553 | 1.40E+06 | 1.55E+06 | 1.77E+06 | 1.78E+06 | 1.68E+06 |
| **AVG** | **556** | **1.45E+06** | **1.62E+06** | **1.72E+06** | **1.71E+06** | **1.57E+06** |
| **SD** | **0.99** | **3.01** | **4.32** | **2.92** | **6.11** | **7.69** |

**2. Effect of mRNA per well on transfection**

**Table S3:** Microscopic images showing transfection efficiency of EGFP-mRNA in 293T and A549 cells at various doses and ratios.

| **Cell line** | **293t** | | |
|---|---|---|---|
| **Dose (ng/well)** | **50** | **100** | **200** |
| Ratio (1:0.5) | 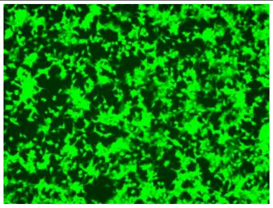 | 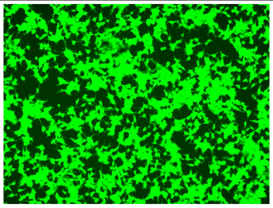 | 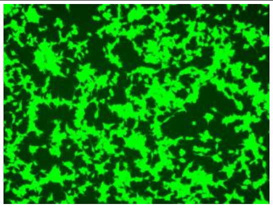 |
| Ratio (1:1) | 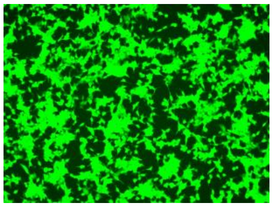 | 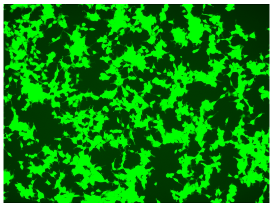 | 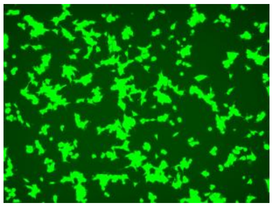 |
| Blank | 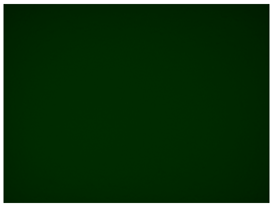 | | |
| **Cell line** | **A549** | | |
| **Dose (ng/well)** | **50** | **100** | **200** |
| Ratio (1:0.5) | 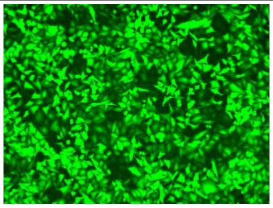 | 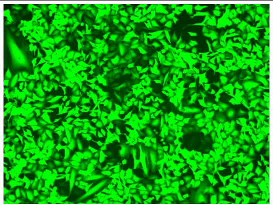 | 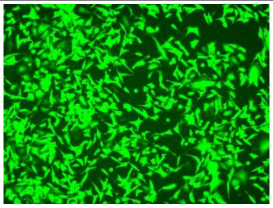 |
| Ratio (1:1) | 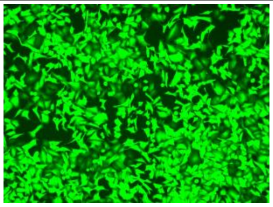 | 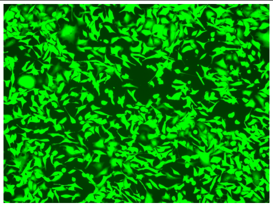 | 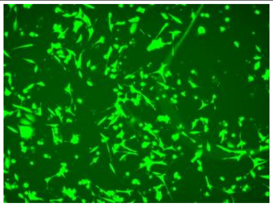 |
| Blank | 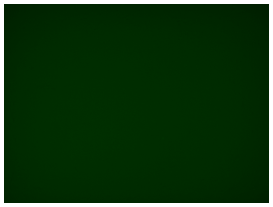 | | |

**Table S4:** Flow cytometry data comparing transfection efficiency in 293T and A549 cells at different doses of EGFP-mRNA.

| **Cell line** | **293t** | **A549** |
|---|---|---|

| **Dose (ng/well)** | **50** | **100** | **200** | **Dose (ng/well)** | **50** | **100** | **200** |
|---|---|---|---|---|---|---|---|
| Ratio 1:0.5 | 8.43E+05 | 1.21E+06 | 1.23E+06 | Ratio (1:0.5) | 5.32E+05 | 1.02E+06 | 1.34E+06 |
| | 7.89E+05 | 1.09E+06 | 1.18E+06 | | 6.44E+05 | 1.08E+06 | 1.01E+06 |
| | 9.22E+05 | 1.19E+06 | 1.08E+06 | | 6.67E+05 | 8.50E+05 | 1.75E+06 |
| | 8.61E+05 | 1.26E+06 | 1.19E+06 | | 4.36E+05 | 1.05E+06 | 1.24E+06 |
| **AVG** | **8.54E+05** | **1.19E+06** | **1.17E+06** | **AVG** | **5.70E+05** | **1.00E+06** | **1.34E+06** |
| Ratio 1:1 | 9.91E+05 | 8.28E+05 | 5.76E+05 | Ratio (1:1) | 9.28E+05 | 1.42E+06 | 1.15E+06 |
| | 1.33E+06 | 1.19E+06 | 4.90E+05 | | 1.27E+06 | 1.57E+06 | 1.02E+06 |
| | 1.07E+06 | 1.19E+06 | 9.09E+05 | | 1.21E+06 | 1.62E+06 | 8.31E+05 |
| | 9.90E+05 | 8.96E+05 | 6.82E+05 | | 1.14E+06 | 1.88E+06 | 1.16E+06 |
| **AVG** | **1.09E+06** | **1.03E+06** | **6.64E+05** | **AVG** | **1.14E+06** | **1.62E+06** | **1.04E+06** |
| blank | 5.78E+02 | | | blank | 5.78E+02 | | |
| | 6.14E+02 | | | | 5.78E+02 | | |
| | 5.06E+02 | | | | 5.78E+02 | | |
| | 5.06E+02 | | | | 3.25E+02 | | |
| **AVG** | **5.51E+02** | | | **AVG** | **5.15E+02** | | |

3. **Effect of mRNA and LNP ratio on transfection**

**Table S5:** Particle size and polydispersity index (PDI) of LNP-RNA complexes formed at different mRNA/LNP ratios.

| **ratio** | **1:0.5** | **1:0.75** | **1:1** | **1:1.25** | **1:1.5** |
|---|---|---|---|---|---|
| Size (nm) | 181.2±3.329 | 171.9±2.335 | 168.0±3.305 | 163.2±3.175 | 162.6±2.007 |
| PDI | 0.130±0.012 | 0.140±0.018 | 0.118±0.033 | 0.132±0.014 | 0.144±0.012 |

**Table S6:** Microscopic images showing the impact of varying mRNA/LNP ratios on transfection.

| Ratio | blank | 1:0.5 | 1:0.75 | 1:1 | 1:1.25 | 1:1.5 |
|---|---|---|---|---|---|---|
| | 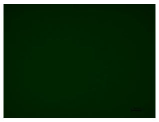 | 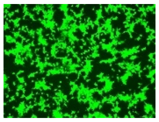 | 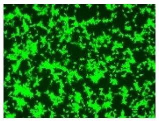 | 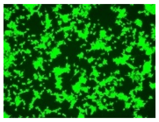 | 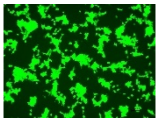 | 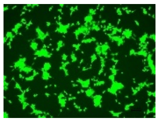 |

**Table S7:** Flow cytometry data displaying the transfection efficiency at different mRNA/LNP ratios.

| Ratio | Blank | 1:0.5 | 1:0.75 | 1:1 | 1:1.25 | 1:1.5 |
|---|---|---|---|---|---|---|
| 1 | 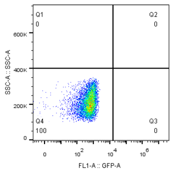 | 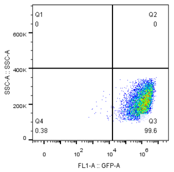 | 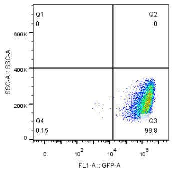 | 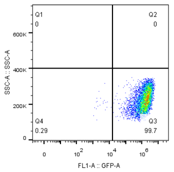 | 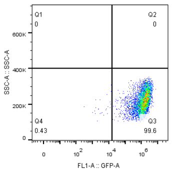 | 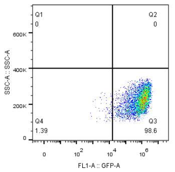 |
| 2 | 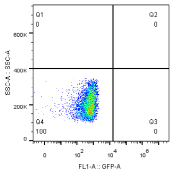 | 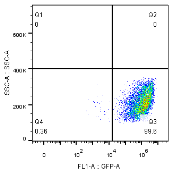 | 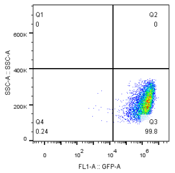 | 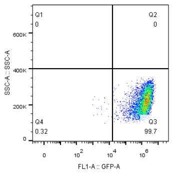 | 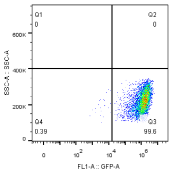 | 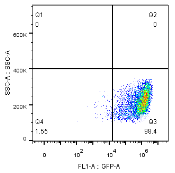 |
| 3 | 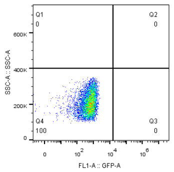 | 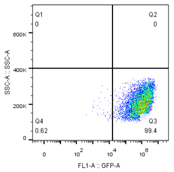 | 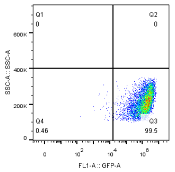 | 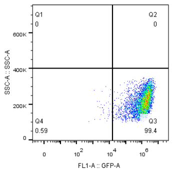 | 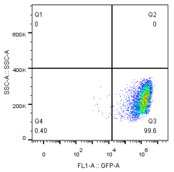 | 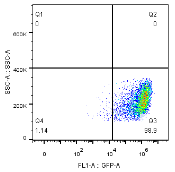 |
| Data 1 | 562 | 1.37E6 | 1.79E6 | 1.69E6 | 1.45E6 | 1.03E6 |
| Data 2 | 552 | 1.42E6 | 1.76E6 | 1.62E6 | 1.42E6 | 1.10E6 |
| Data 3 | 553 | 1.26E6 | 1.68E6 | 1.55E6 | 1.42E6 | 1.11E6 |
| **AVG** | **556** | **1.35E+06** | **1.74E+06** | **1.62E+06** | **1.43E+06** | **1.08E+06** |
| **SD** | **0.99** | **6.06** | **3.26** | **4.32** | **1.21** | **4.04** |

**4. Mixing techniques**

**Table S8:** Particle size and PDI of LNP-RNA complexes formed using different mixing techniques at varying ratios.

| Ratio | Mixing technique | Size (nm) | PDI |
|---|---|---|---|
| 1:0.5 | Pipetting | 138.4±2.312 | 0.168±0.048 |
| | Vortex 3s | 144.2±1.429 | 0.179±0.025 |
| | Vortex 10s | 159.5±3.164 | 0.210±0.013 |
| 1:1 | Pipetting | 190.4±2.012 | 0.165±0.048 |
| | Vortex 3s | 203.9±1.662 | 0.125±0.006 |
| | Vortex 10s | 209.2±3.955 | 0.152±0.016 |

**Table S9:** Microscopic images illustrating the effect of different mixing techniques on LNP-RNA complex formation.

| Ratio | Pipetting | Vortex 3s | Vortex10s |
|---|---|---|---|
| 1:0.5 | 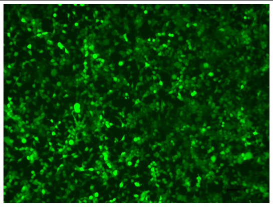 | 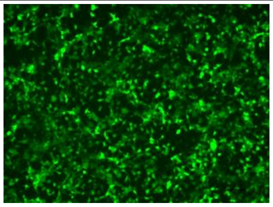 | 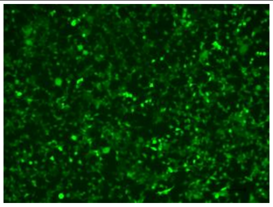 |
| 1:1 | 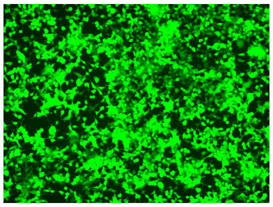 | 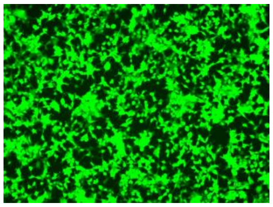 | 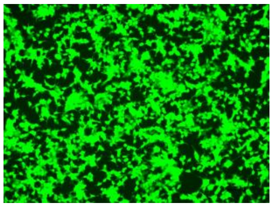 |
| blank | 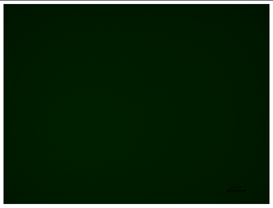 | | |

**Table S10:** Flow cytometry data showing transfection efficiency at different mRNA/LNP ratios and mixing techniques.

| **mRNA/LNP ratio = 1:1** | | | |
|---|---|---|---|
| Mixing | Pipetting | Vortex 3s | Vortex 10s |

| | | | |
|---|---|---|---|
| 1 | 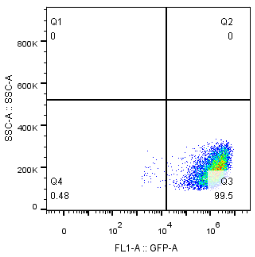 | 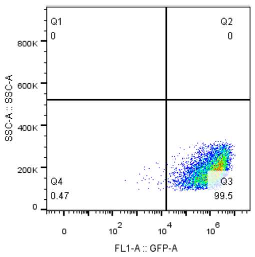 | 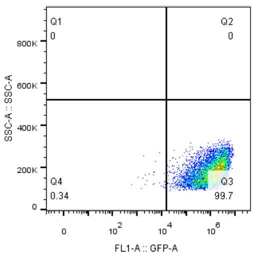 |
| 2 | 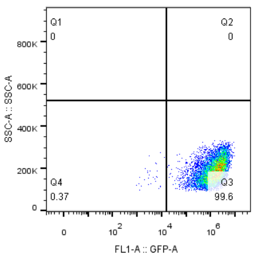 | 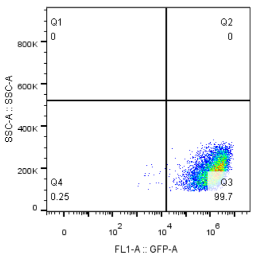 | 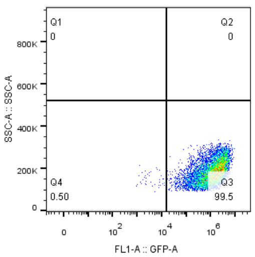 |
| 3 | 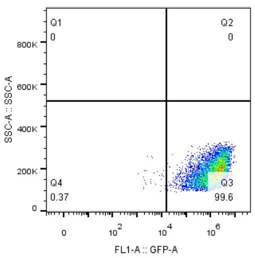 | 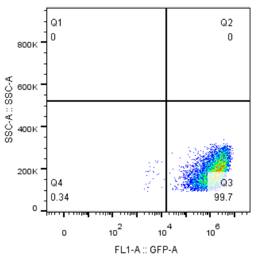 | 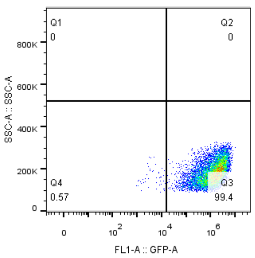 |
| Data1 | 1.73E+06 | 1.53E+06 | 1.50E+06 |
| Data 2 | 1.71E+06 | 1.65E+06 | 1.58E+06 |
| Data 3 | 1.65E+06 | 1.73E+06 | 1.61E+06 |
| **AVG** | **1.70E+06** | **1.64E+06** | **1.56E+06** |
| **SD** | **2.45** | **6.15** | **3.64** |

| **mRNA/LNP ratio = 1:0.5** | | | | |
|---|---|---|---|---|
| Mixing | Pipetting | Vortex 3s | Vortex 10s | blank |
| 1 | 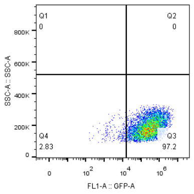 | 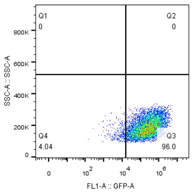 | 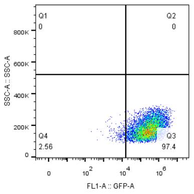 | 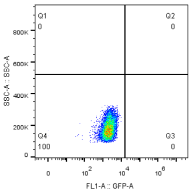 |

| | | | | |
|---|---|---|---|---|
| 2 | 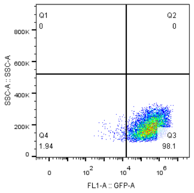 | 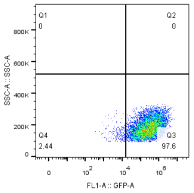 | 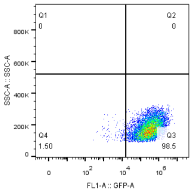 | |
| 3 | 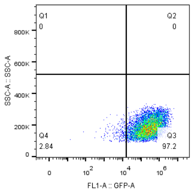 | 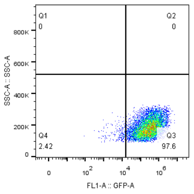 | 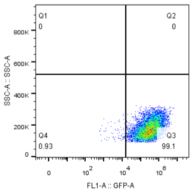 | |
| Data 1 | 4.60E+05 | 4.17E+05 | 4.61E+05 | 2197 |
| Data 2 | 4.89E+05 | 4.56E+05 | 4.59E+05 | |
| Data 3 | 4.31E+05 | 4.50E+05 | 4.66E+05 | |
| **AVG** | **4.60E+05** | **4.41E+05** | **4.62E+05** | |
| **SD** | **6.30** | **4.76** | **0.78** | |

### 5. Stability of mRNA-LNP complexes

**Table S11:** Microscopic images showing the stability of mRNA-LNP complexes over time.

| Time(h) | 0 | 0.5 | 1 | 3 | 8 | 24 |
|---|---|---|---|---|---|---|
| 1:1 | 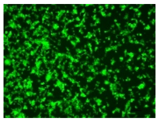 | 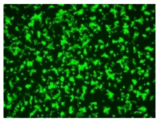 | 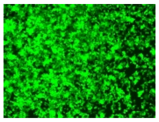 | 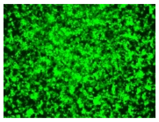 | 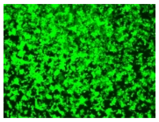 | 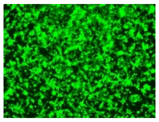 |
| 1:0.5 | 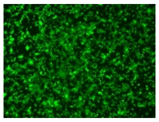 | 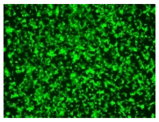 | / | 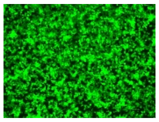 | 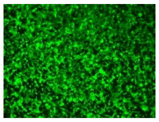 | 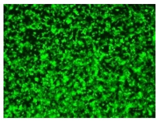 |
| Blank | 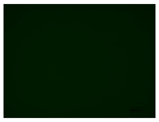 | / | / | / | / | / |

**Table S12:** Flow cytometry data assessing transfection efficiency over time for different mRNA/LNP ratios.

| **mRNA/LNP ratio = 1:1** | | | | | | | |
|---|---|---|---|---|---|---|---|
| Time (h) | 0 | 0.5 | 1 | 3 | 8 | 24 | Blank |
| 1 | 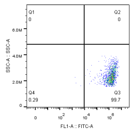 | 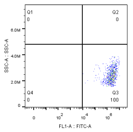 | 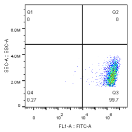 | 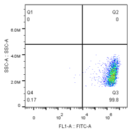 | 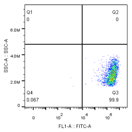 | 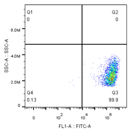 | 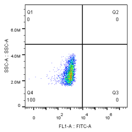 |
| 2 | 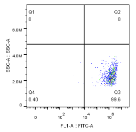 | 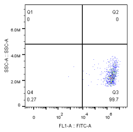 | 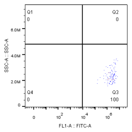 | 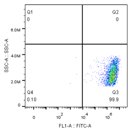 | 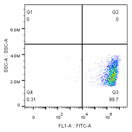 | 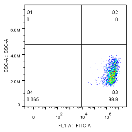 | |
| 3 | 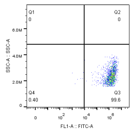 | 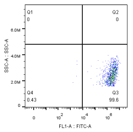 | 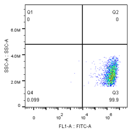 | 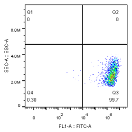 | 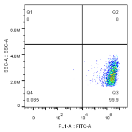 | 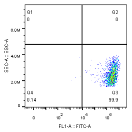 | |
| Data 1 | 1.62E+06 | 2.52E+06 | 2.63E+06 | 2.77E+06 | 2.63E+06 | 2.60E+06 | 945 |
| Data 2 | 1.63E+06 | 2.59E+06 | 2.40E+06 | 2.75E+06 | 2.66E+06 | 3.08E+06 | |
| Data 3 | 1.61E+06 | 2.61E+06 | 2.36E+06 | 2.62E+06 | 2.65E+06 | 2.99E+06 | |
| **AVG** | **1.62E+06** | **2.57E+06** | **2.46E+06** | **2.71E+06** | **2.65E+06** | **2.89E+06** | |
| **SD** | **0.62** | **1.84** | **5.92** | **3.00** | **0.58** | **8.83** | |

| **mRNA/LNP ratio = 1:0.5** | | | | | | | |
|---|---|---|---|---|---|---|---|
| Time (h) | 0 | 0.5 | 1 | 3 | 8 | 24 | Blank |
| 1 | 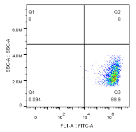 | 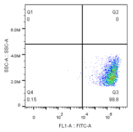 | 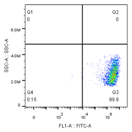 | 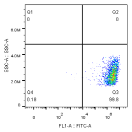 | 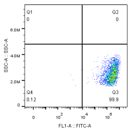 | 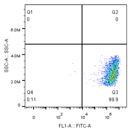 | 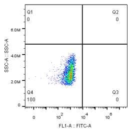 |

| | | | | | | | |
|---|---|---|---|---|---|---|---|
| 2 | 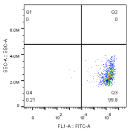 | 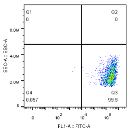 | 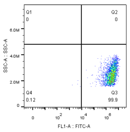 | 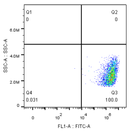 | 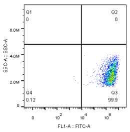 | 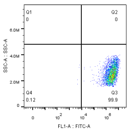 | |
| 3 | 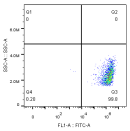 | 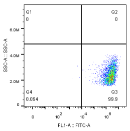 | 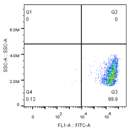 | 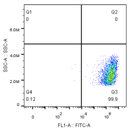 | 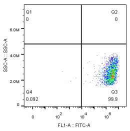 | 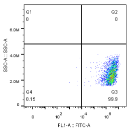 | |
| Data 1 | 1.65E+06 | 2.46E+06 | 3.11E+06 | 2.86E+06 | 2.80E+06 | 3.01E+06 | 945 |
| Data 2 | 1.78E+06 | 2.50E+06 | 3.01E+06 | 2.64E+06 | 2.75E+06 | 3.14E+06 | |
| Data 3 | 1.71E+06 | 2.48E+06 | 2.95E+06 | 2.42E+06 | 2.86E+06 | 2.90E+06 | |
| **AVG** | **1.71E+06** | **2.48E+06** | **3.02E+06** | **2.64E+06** | **2.80E+06** | **3.02E+06** | |
| **SD** | **3.80** | **0.81** | **2.67** | **8.33** | **1.96** | **3.98** | |

### 6. Influence of cμlture medium on transfection

**Table S13:** Microscopic images showing the effect of culture medium composition on transfection efficiency.

| Ratio | serum(+)/antibiotics(-) | serum(-)/antibiotics(-) | serum(+)/antibiotics(+) |
|---|---|---|---|
| 1:0.5 | 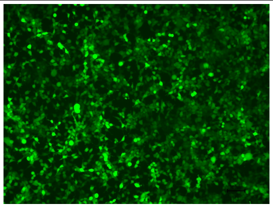 | 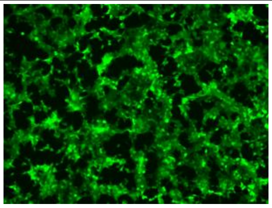 | 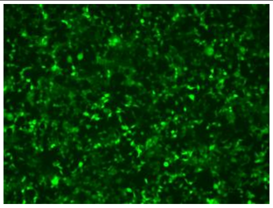 |
| 1:1 | 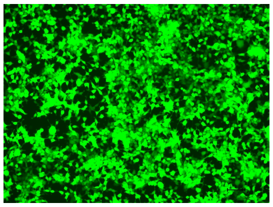 | 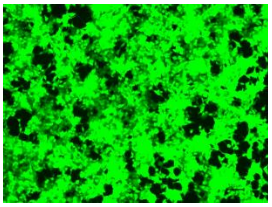 | 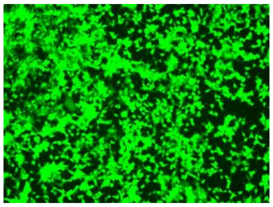 |

**Table S14:** Flow cytometry data comparing transfection efficiency across various culture media conditions.

<table>
<tr><th colspan="4">mRNA/LNP ratio = 1:1</th></tr>
<tr><th></th><th>serum(+)/antibiotics(-)</th><th>serum(-)/antibiotics(-)</th><th>serum(+)/antibiotics(+)</th></tr>
<tr><td>1</td><td>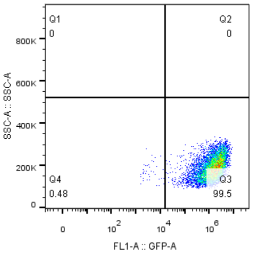
</td><td>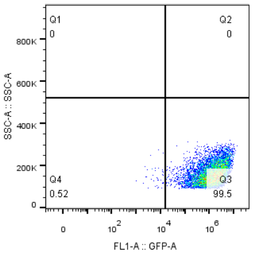
</td><td>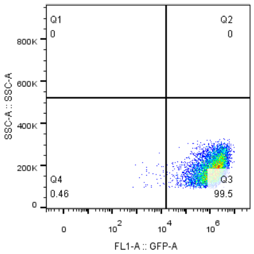
</td></tr>
<tr><td>2</td><td>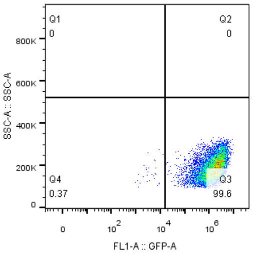
</td><td>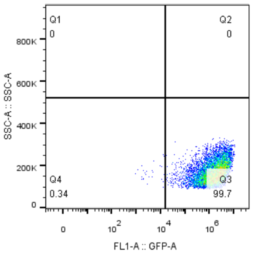
</td><td>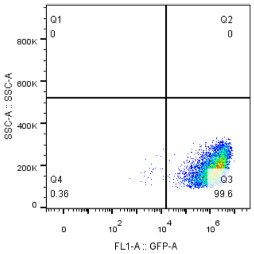
</td></tr>
<tr><td>3</td><td>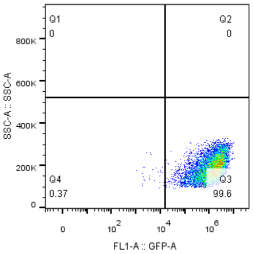
</td><td>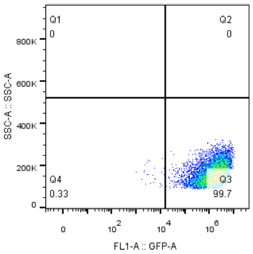
</td><td>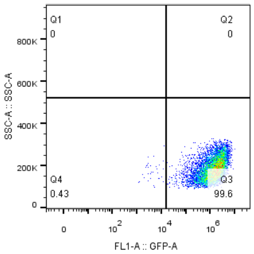
</td></tr>
</table>

| | | | |
|---|---|---|---|
| Data 1 | 1.73E+06 | 1.85E+06 | 1.53E+06 |
| Data 2 | 1.71E+06 | 2.04E+06 | 1.74E+6 |
| Data 3 | 1.65E+06 | 1.89E+06 | 1.65E+06 |
| **AVG** | **1.70E+06** | **1.93E+06** | **1.64E+06** |
| **SD** | **2.45** | **5.20** | **6.42** |

<table>
<tr><td colspan="4">mRNA/LNP ratio = 1:0.5</td></tr>
<tr><td></td><td>serum(+)/antibiotics(-)</td><td>serum(-)/antibiotics(-)</td><td>serum(+)/antibiotics(+)</td></tr>
<tr><td>1</td><td>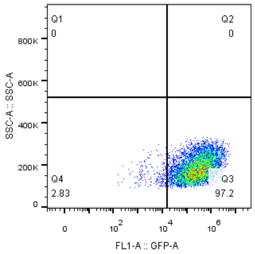</td><td>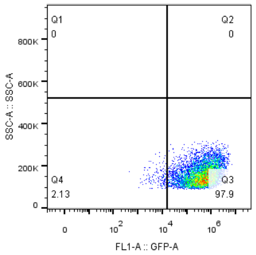</td><td>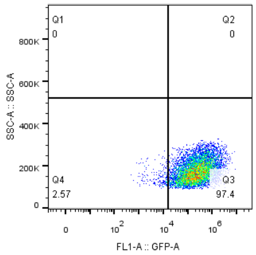</td></tr>
<tr><td>2</td><td>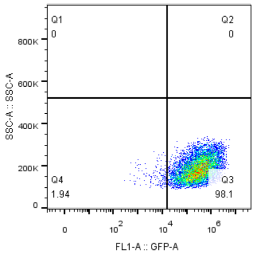</td><td>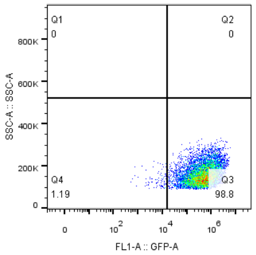</td><td>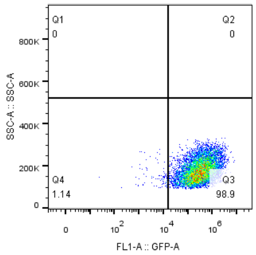</td></tr>
<tr><td>3</td><td>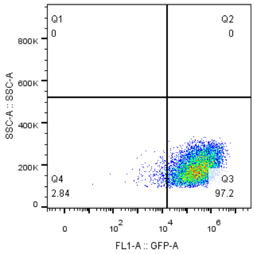</td><td>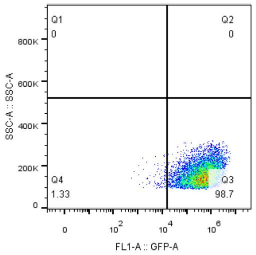</td><td>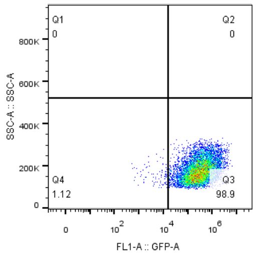</td></tr>
<tr><td>Data 1</td><td>4.60E+05</td><td>6.62E+05</td><td>4.04E+05</td></tr>
<tr><td>Data 2</td><td>4.89E+05</td><td>6.73E+05</td><td>4.85E+05</td></tr>
<tr><td>Data 3</td><td>4.31E+05</td><td>6.55E+05</td><td>4.78E+05</td></tr>
<tr><td>AVG</td><td>4.60E+05</td><td>6.63E+05</td><td>4.56E+05</td></tr>
<tr><td>SD</td><td>6.30</td><td>1.37</td><td>9.85</td></tr>
</table>

**7. Fluorescent intensity changes with time**

**Table S15:** Flow cytometry data showing changes in fluorescent intensity over time for different mRNA/LNP ratios.

| **Cell line** | **293t Luc-mRNA** | | | | |
|---|---|---|---|---|---|
| **Time** | **6h** | **12h** | **24** | **48** | **72** |
| SD 1:0.5 | 5.54E+05 | 1.52E+06 | 1.21E+06 | 8.38E+05 | 1.24E+05 |
| | 2.78E+05 | 1.35E+06 | 1.09E+06 | 6.00E+05 | 1.85E+05 |
| | 3.39E+05 | 7.84E+05 | 1.19E+06 | 6.71E+05 | 2.00E+05 |
| | 4.98E+05 | 1.37E+06 | 1.26E+06 | 5.69E+05 | 1.75E+05 |
| **AVG** | **4.17E+05** | **1.26E+06** | **1.19E+06** | **6.70E+05** | **1.71E+05** |
| SD 1:1 | 8.12E+05 | 1.30E+06 | 8.28E+05 | 4.68E+05 | 7.43E+04 |
| | 6.51E+05 | 1.16E+06 | 1.19E+06 | 5.97E+05 | 1.43E+05 |
| | 8.04E+05 | 1.27E+06 | 1.19E+06 | 3.48E+05 | 1.23E+05 |
| | 5.23E+05 | 7.64E+05 | 8.96E+05 | 3.60E+05 | 9.34E+04 |
| **AVG** | **6.97E+05** | **1.12E+06** | **1.03E+06** | **4.43E+05** | **1.08E+05** |
| blank | 5.00E+02 | 1.00E+03 | 5.78E+02 | 8.96E+02 | 3.11E+02 |
| | 2.08E+02 | 1.16E+03 | 6.14E+02 | 1.02E+03 | 5.84E+02 |
| | 2.50E+02 | 6.94E+02 | 5.06E+02 | 8.55E+02 | 5.45E+02 |
| | 3.33E+02 | 5.79E+02 | 5.06E+02 | 7.74E+02 | 1.56E+02 |
| **AVG** | **3.23E+02** | **8.58E+02** | **5.51E+02** | **8.86E+02** | **3.99E+02** |

| **Cell lines** | **A549 Luc-mRNA** | | | | |
|---|---|---|---|---|---|
| **Time** | **6h** | **12h** | **24** | **48** | **72** |
| SD 1:0.5 | 3.97E+05 | 7.87E+05 | 1.02E+06 | 3.69E+05 | 5.75E+04 |
| | 2.04E+05 | 1.11E+06 | 1.08E+06 | 4.58E+05 | 5.11E+04 |
| | 1.96E+05 | 8.51E+05 | 8.50E+05 | 5.22E+05 | 8.40E+04 |
| | 3.62E+05 | 1.10E+06 | 1.05E+06 | 3.48E+05 | 9.16E+04 |
| **AVG** | **2.90E+05** | **9.61E+05** | **1.00E+06** | **4.24E+05** | **7.10E+04** |
| SD 1:1 | 4.81E+05 | 1.15E+06 | 1.42E+06 | 4.44E+05 | 9.74E+04 |
| | 6.63E+05 | 1.49E+06 | 1.57E+06 | 2.90E+05 | 8.66E+04 |
| | 3.27E+05 | 1.42E+06 | 1.62E+06 | 2.95E+05 | 8.01E+04 |
| | 5.56E+05 | 1.91E+06 | 1.88E+06 | 5.65E+05 | 1.25E+05 |
| **AVG** | **5.07E+05** | **1.49E+06** | **1.62E+06** | **3.98E+05** | **9.73E+04** |
| blank | 3.75E+02 | 4.24E+02 | 5.78E+02 | 7.74E+02 | 2.72E+02 |
| | 2.50E+02 | 7.33E+02 | 5.78E+02 | 5.29E+02 | 2.34E+02 |
| | 2.08E+02 | 6.94E+02 | 5.78E+02 | 7.33E+02 | 2.72E+02 |
| | 2.92E+02 | 7.33E+02 | 3.25E+02 | 5.70E+02 | 2.34E+02 |
| **AVG** | **2.81E+02** | **6.46E+02** | **5.15E+02** | **6.52E+02** | **2.53E+02** |

**Table S16**: Microscopic images showing transfection efficiency over time in 293T and A549 cells.

| Cell line | 293t | | | | | |
|---|---|---|---|---|---|---|
| **Time** | **6h** | **12h** | **24** | **48h** | **72h** | **7d** |
| SD1:0.5 | 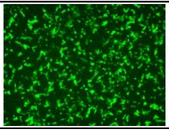 | 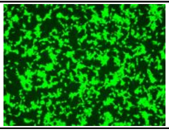 | 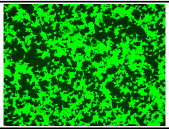 | 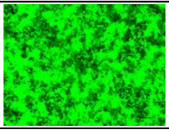 | 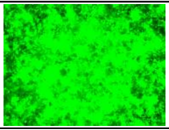 | 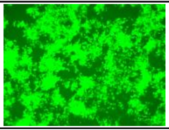 |
| SD1:1 | 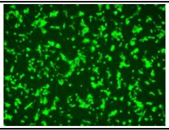 | 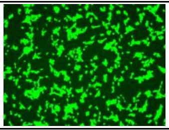 | 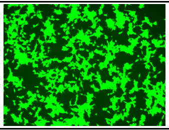 | 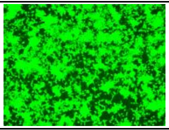 | 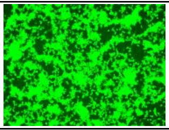 | 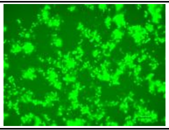 |
| blank | 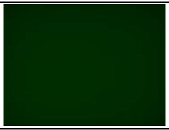 | 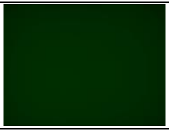 | 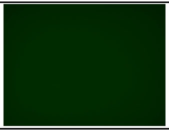 | 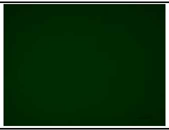 | 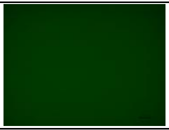 | 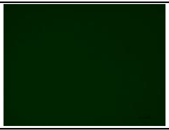 |

| Cell line | A549 | | | | | |
|---|---|---|---|---|---|---|
| **Time** | **6h** | **12h** | **24h** | **48h** | **72h** | **7d** |
| SD1:0.5 | 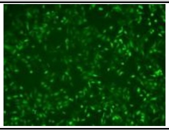 | 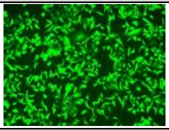 | 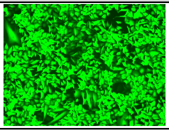 | 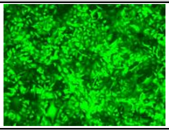 | 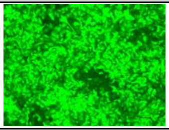 | 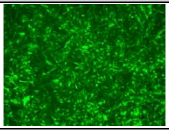 |
| SD1:1 | 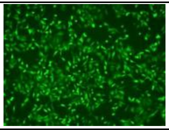 | 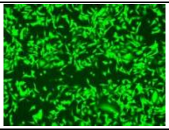 | 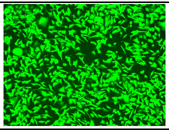 | 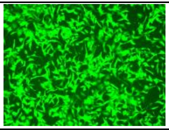 | 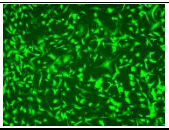 | 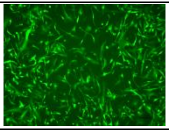 |
| blank | 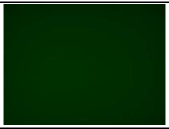 | 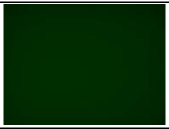 | 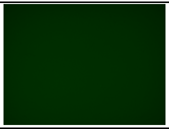 | 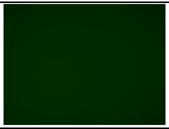 | 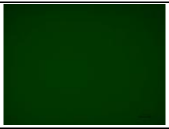 | 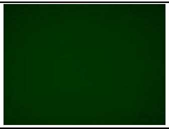 |

8. **RNA types**

**Table S17:** Particle size and PDI of LNP complexes containing different RNA types (mRNA, circRNA, saRNA).

| Ratio | RNA type | Size (nm) | PDI |
|---|---|---|---|
| 1:0.5 | EGFP-mRNA | 138.4±2.312 | 0.168±0.048 |
| | EGFP-circRNA | 179.4±1.343 | 0.174±0.013 |
| | EGFP-saRNA | 251.8±2.155 | 0.317±0.025 |
| 1:1 | EGFP-mRNA | 190.4±2.012 | 0.165±0.048 |
| | EGFP-circRNA | 175.8±1.601 | 0.156±0.019 |
| | EGFP-saRNA | 191.8±2.829 | 0.214±0.028 |

**Table S18**: Microscopic images comparing the transfection efficiency of mRNA, circRNA, and saRNA.

| Ratio | circRNA | saRNA | mRNA |
|---|---|---|---|
| 1:0.5 | 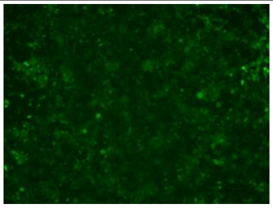 | 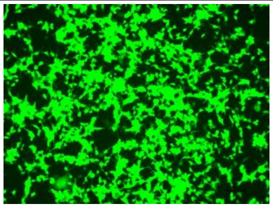 | 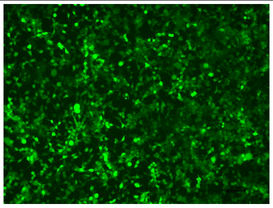 |
| 1:1 | 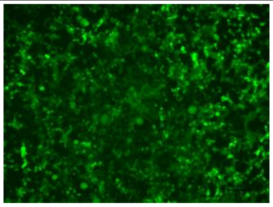 | 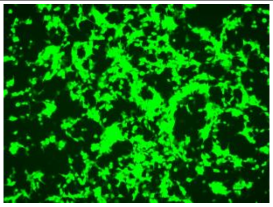 | 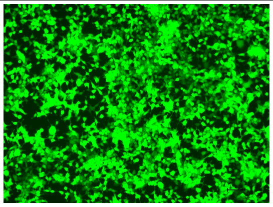 |
| Blank | | | 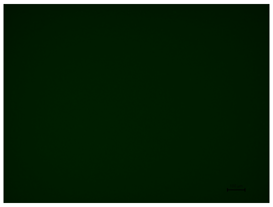 |

**Table S19:** Flow cytometry data comparing the transfection efficiency of mRNA, circRNA, and saRNA at varying ratios.

| **mRNA/LNP ratio = 1:1** | | | |
|---|---|---|---|
| | CircRNA | SaRNA | mRNA |
| 1 | 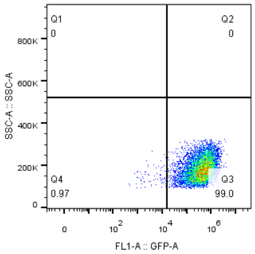  | 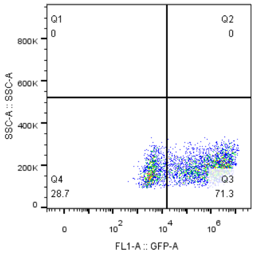  | 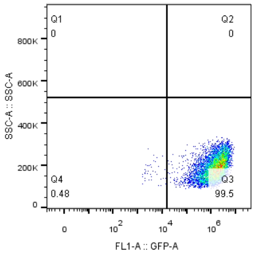  |

| | | | |
|---|---|---|---|
| 2 | 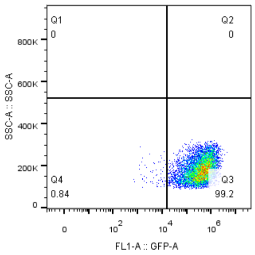 | 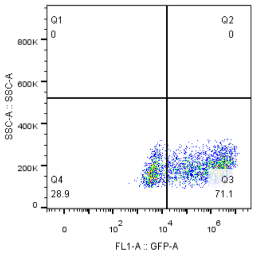 | 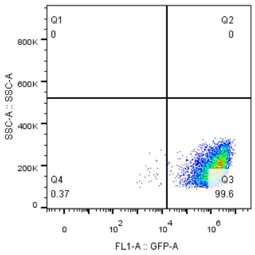 |
| 3 | 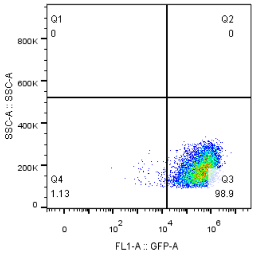 | 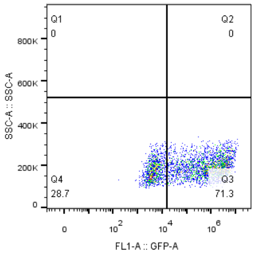 | 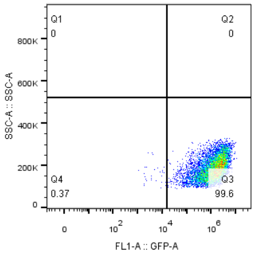 |
| Data 1 | 5.04E+05 | 1.24E+06 | 1.73E+06 |
| Data 2 | 4.91E+05 | 1.20E+06 | 1.71E+06 |
| Data 3 | 4.93E+05 | 1.15E+06 | 1.65E+06 |
| **AVG** | **4.96E+05** | **1.20E+06** | **1.70E+06** |
| **SD** | **1.41** | **3.77** | **2.45** |

| **mRNA/LNP ratio = 1:0.5** | | | |
|---|---|---|---|
| | CircRNA | SaRNA | mRNA |
| 1 | 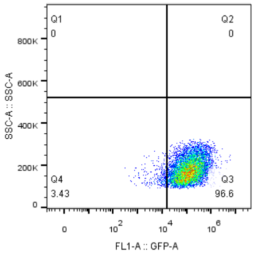 | 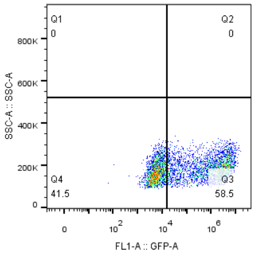 | 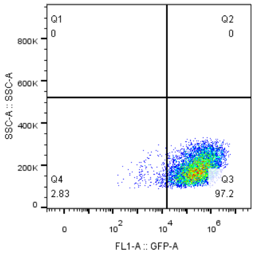 |
| 2 | 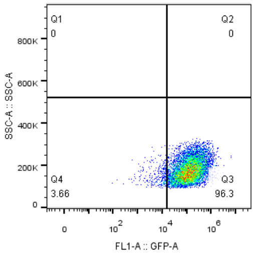 | 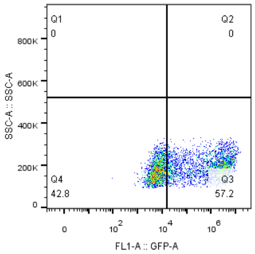 | 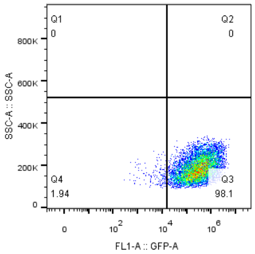 |

| | | | |
|---|---|---|---|
| 3 | 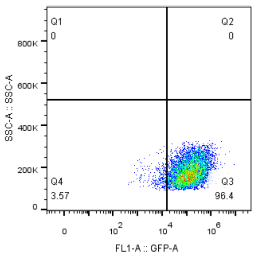 | 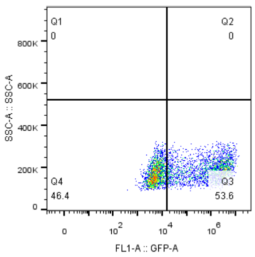 | 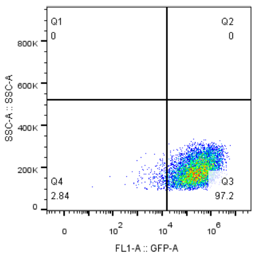 |
| 1 | 1.93E+05 | 1.15E+06 | 4.60E+05 |
| 2 | 1.92E+05 | 1.12E+06 | 4.89E+05 |
| 3 | 1.97E+05 | 9.74E+05 | 4.31E+05 |
| **AVG** | **1.94E+05** | **1.08E+06** | **4.60E+05** |
| **SD** | **1.36** | **8.71** | **6.30** |

#### 9. Transfection efficiency in different cell lines

**Table S20:** Microscopic images showing transfection efficiency in different cell lines (293T, Jurkat, THP-1).

| | 1:0.5 | 1:1 | Blank |
|---|---|---|---|
| 293t | 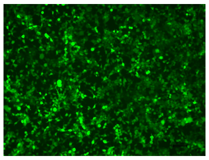 | 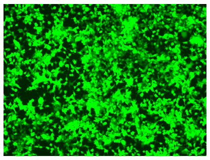 | 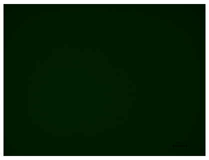 |
| Jurkat | 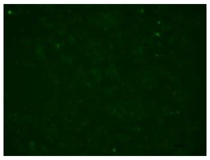 | 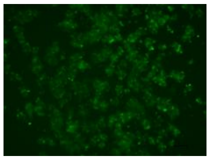 | 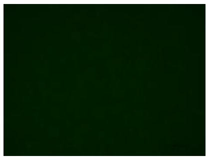 |
| THP-1 | 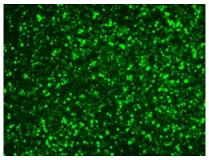 | 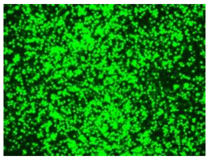 | 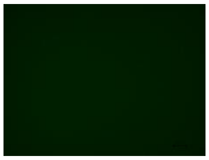 |

**Table S21:** Flow cytometry data comparing transfection efficiency in 293T, Jurkat, and THP-1 cell lines.

| **mRNA/LNP ratio = 1:1** | | | |
|---|---|---|---|
| | 293t | Jurkat | THP-1 |
| 1 | 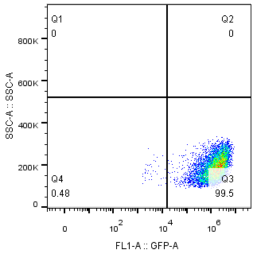  | 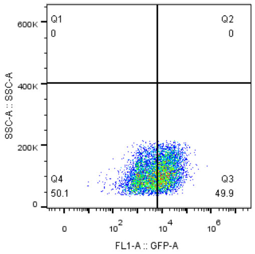  | 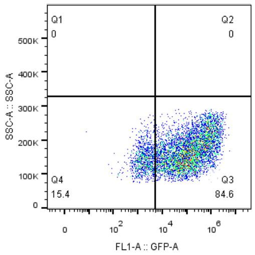  |
| 2 | 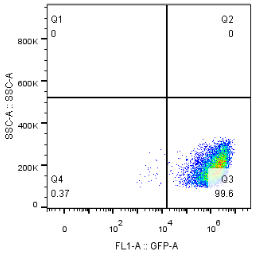  | 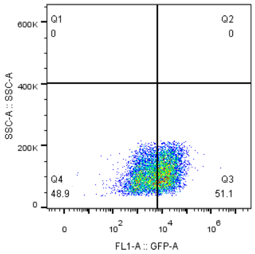  | 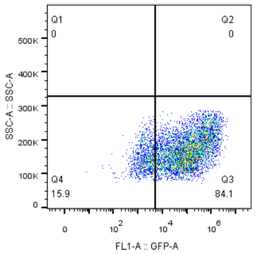  |

| | | | |
|---|---|---|---|
| 3 | 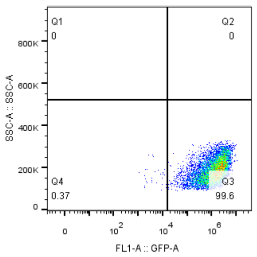 | 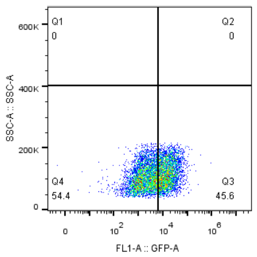 | 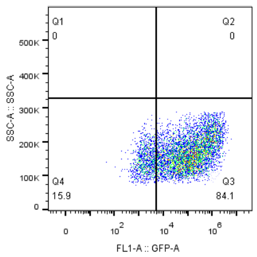 |
| 1 | 1.73E+06 | 1.21E+04 | 3.17E+05 |
| 2 | 1.71E+06 | 1.23E+04 | 2.78E+05 |
| 3 | 1.65E+06 | 1.00E+04 | 3.15E+05 |
| **AVG** | **1.70E+06** | **1.15E+04** | **3.03E+05** |
| **SD** | **2.45** | **11.11** | **7.24** |

| **mRNA/LNP ratio = 1:0.5** | | | |
|---|---|---|---|
| | 293t | Jurkat | THP-1 |
| 1 | 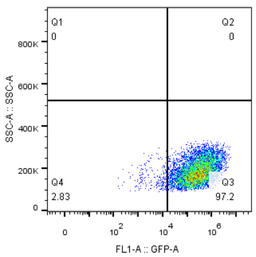 | 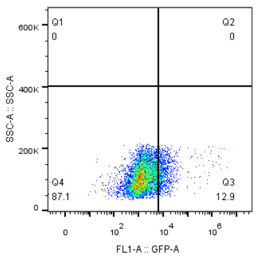 | 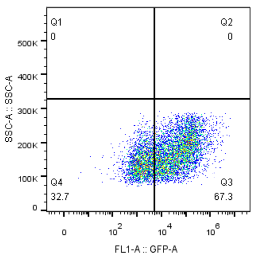 |
| 2 | 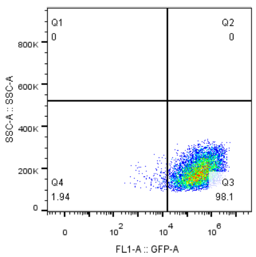 | 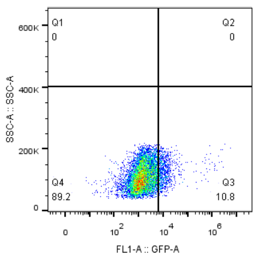 | 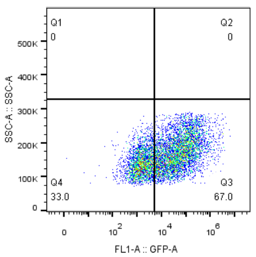 |
| 3 | 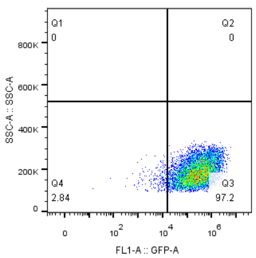 | 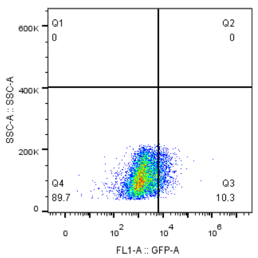 | 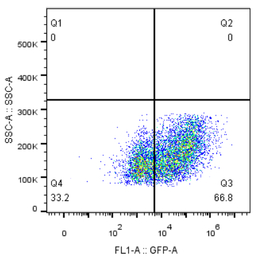 |
| 1 | 4.60E+05 | 7.81E+03 | 9.60E+04 |

| | | | |
|---|---|---|---|
| 2 | 4.89E+05 | 8.09E+03 | 9.84E+04 |
| 3 | 4.31E+05 | 3.13E+03 | 8.94E+04 |
| **AVG** | **4.60E+05** | **6.34E+03** | **9.46E+04** |
| **SD** | **6.30** | **43.93** | **4.93** |
